# Solid-Phase Synthesis of ProTide Fluorogenic Probes Enables Systematic Profiling of Carboxypeptidase Activity

**DOI:** 10.64898/2026.03.10.710453

**Authors:** Toru Komatsu, Mayano Minoda, Takako Uchida, Momoka Hata, Shunsuke Kanai, Hideto Hiraide, Yu Kagami, Kazufumi Honda, Yasuteru Urano

**Author notes:** **Corresponding Author:** Toru Komatsu, Yasuteru Urano.

## Abstract

Carboxypeptidases play diverse roles in physiological and pathological processes, yet comprehensive analysis of their activities in complex biological samples remains challenging. Here we report a solid-phase synthesis strategy for fluorogenic ProTide-based probes that enables systematic profiling of carboxypeptidase activities based on defined C-terminal amino acid motifs. By modular synthesis of dipeptide–fluorophore conjugates, we generated a focused probe set that revealed distinct substrate preferences among carboxypeptidases, including carboxypeptidase A and B family enzymes. Integration of these probes with a single-molecule enzyme activity assay allowed ultrasensitive detection of circulating carboxypeptidase activities in human blood samples. Application of this platform to clinical specimens demonstrated that specific carboxypeptidase activities are elevated in patients with pancreatic cancer compared with healthy controls, whereas closely related enzymes showed limited diagnostic value. These results establish a scalable chemical strategy for activity-based profiling of exopeptidases and highlight circulating carboxypeptidase activity as a functional enzymatic signature associated with pancreatic cancer.

## Introduction

Enzyme activity is a central determinant of physiological and pathological processes, and its quantitative monitoring is fundamental to biochemical analysis, drug discovery, and disease diagnostics^1–3^. Fluorogenic probes serve as one of the most powerful tools for directly monitoring enzyme activity due to their high sensitivity, operational simplicity, and compatibility with various experimental platforms for *in vitro, in cellulo*, and *in vivo* assays.

Despite their widespread utility, the design of fluorogenic probes that achieve selectivity among closely related enzyme isoforms remains a major challenge. Subtle differences in substrate preference, catalytic mechanisms, and physiological localization often require fine-tuned substrate structures to distinguish individual enzyme activities in complex biological environments. This challenge is particularly pronounced for enzyme families with multiple members exhibiting overlapping substrate specificities^2^.

Carboxypeptidases represent a family of exopeptidases that catalyze the removal of C-terminal amino acids from peptide substrates. Nearly 20 human enzymes exhibiting carboxypeptidase-type activities have been identified that regulate diverse biological processes including digestive peptide processing, hormone maturation and extracellular signaling (**Table S1**). While dysregulation of carboxypeptidase activity has been implicated in multiple disease states, systematic biochemical tools to profile their activity with sequence-level selectivity are lacking. In particular, fluorogenic probes capable of distinguishing among carboxypeptidase isoforms in complex biological samples have not been comprehensively developed^4–7^. We have shown that ProTide-based fluorogenic probes provide a promising strategy to address this challenge^5^. In this design, the fluorophore-linked phosphate (or phosphonate), namely ProTide probe reports peptide cleavage through intramolecular cyclization of the newly generated mono-amino acid, resulting in fluorophore release (**Figure 1a**). Due to sequence flexibility, it gives the versatile strategy to monitor dipeptide cleavage reaction at C-terminus, but previous applications of ProTide chemistry have largely relied on a limited number of substrate sequences, leaving the substrate specificity landscape of carboxypeptidases largely unexplored^5^. A major barrier has been the synthetic accessibility of diverse ProTide substrates, as conventional solution-phase synthesis requires multiple protection/deprotection and coupling steps and incurs high material cost.

**Figure 1.**
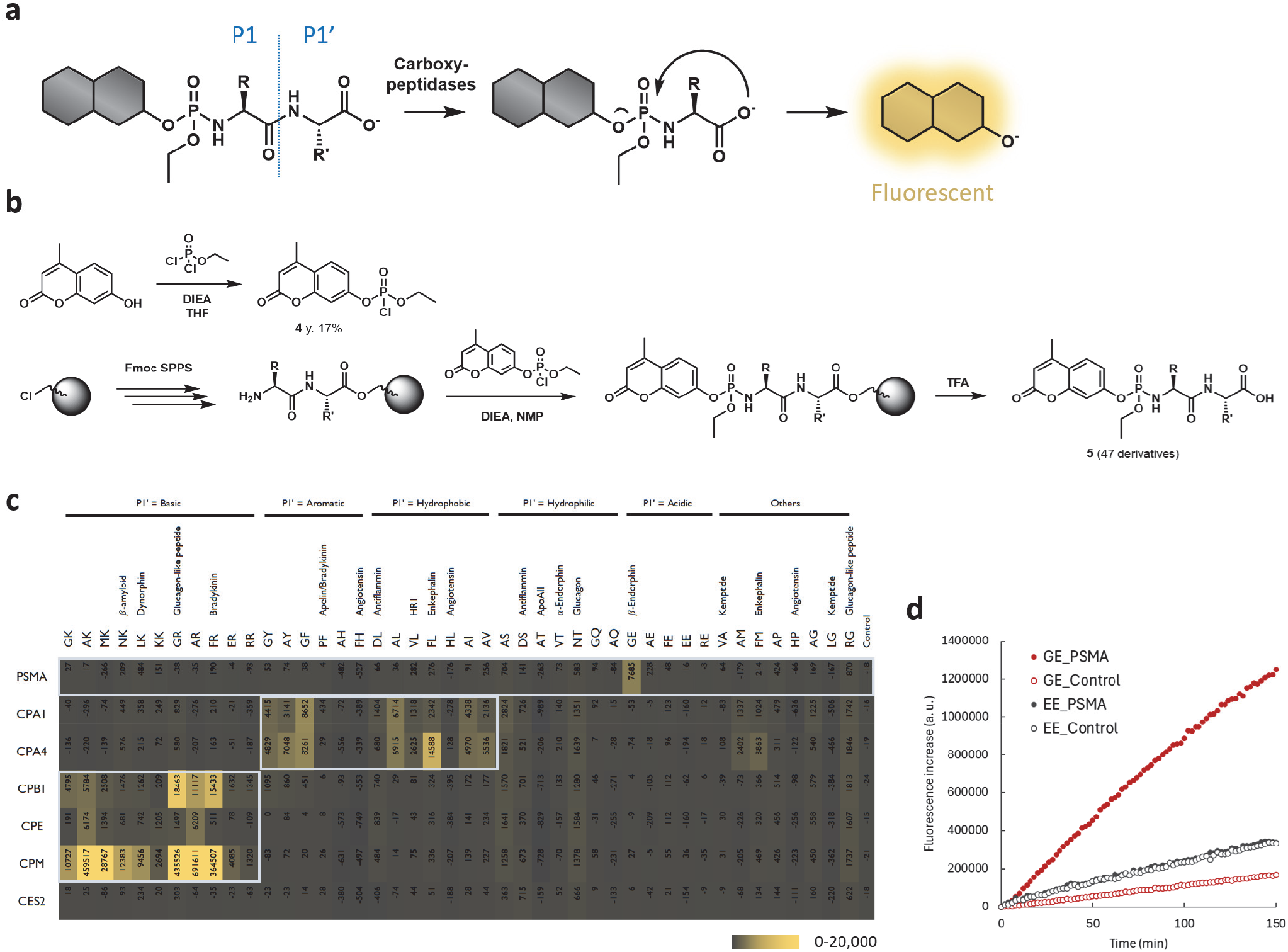
Solid-phase synthesis platform of ProTide probes. (a) Reaction mechanism of ProTide-based probes to monitor carboxypeptidase activities. (b) Synthetic scheme of ProTide-based probe using solid-phase. (c) The result of activity monitoring. The value indicates the fluorescence increase rate (/min) on plate reader. The horizontal entry shows the dipeptide attached to the ProTide, and for the sequences derived from the physiological peptide substrates, their names are also showh. The vertical entry shows the recombinant enzymes. The assay was performed by mixing 4-MU probes (30 μM) and carboxypeptidases (7.5 μg/mL, see Supplementary Information for the details) in Dulbecco’s phosphate buffered saline (DPBS, pH 7.4) containing CHAPS (0.1%) and incubating at 25°C for 150 min. Bar graphs of the results of the representative enzymes are shown in **Figure S2, S3.**(d) Time course change of fluorescence increase (background subtracted) of 4MU-P(Et)-GE (red) and 4MU-P(Et)-EE (black) with or without PSMA (7.5 μg/mL). The condition was same as that of (c).

To overcome these limitations and enable systematic exploration of carboxypeptidase substrate preferences for better substrate designs, we sought to establish a solid-phase synthetic platform for generating ProTide-based fluorogenic probes (**Figure 1b**). This strategy allows rapid and parallel diversification of peptide sequences, enabling comprehensive profiling of enzyme-specific reactivity. By integrating this probe library with biochemical analysis and single-molecule enzyme activity analysis, we sought to uncover isoform-selective substrates and apply them to the detection of disease-associated alterations of enzyme activities in blood samples.

### Solid-phase synthesis of ProTide-based fluorogenic probes

Conventional ProTide synthesis relies on a three-component phosphoramidate-forming reaction in solution, where the stoichiometric balance of reagents is highly sensitive and often difficult to control (**Scheme S1**). These conditions are generally incompatible with solid-phase synthesis, where excess reagents are required to drive reactions to completion. To overcome this challenge, we sought to isolate phosphoro-and phosphono-chloridate intermediates in a stable and purified form, allowing their use in large molar excess during on-resin coupling. We revealed that several phosphorochloridates could be obtained in sufficient purity following chromatographic isolation. One of the examples was the combination of 4-methylumbelliferone (4-MU), a blue-fluorescent fluorophore, and ethyl phosphorochloridate, found to provide sufficient stability for purification and sufficient reactivity to form phosphoramidate on solid phase (**Figure 1b**). It was therefore employed as the basis for library construction. Dipeptide substrates were assembled on 2-chlorotrityl chloride (CTC) resin using Fmoc solid-phase peptide synthesis, after which the N-terminal amine was converted into the ProTide moiety through on-resin reaction with the isolated phosphorochloridate (**Scheme S2**). After the cleavage, most probes were acquired as a pure form (monitored at 320 nm absorbance), so the scheme enabled straightforward diversification of peptide sequences and provided a modular route for probe synthesis.

### Reactivity profiling and structure–activity relationships across carboxypeptidases

The solid-phase platform enabled rapid and standardized preparation of a library of 4-MU–based ProTide probes bearing diverse C-terminal peptide motifs. During library construction, we observed pronounced sequence-dependent differences in probe stability. In particular, probes containing serine or threonine at the P1 position underwent rapid hydrolysis during or after the final deprotection step and were therefore largely unobtainable. This instability is likely attributable to intramolecular nucleophilic attack by the side-chain hydroxyl or carboxyl groups on the electrophilic phosphorus center, resulting in premature fluorophore release.

Excluding these unstable motifs, we obtained a panel of 47 structurally diverse probes suitable for biochemical evaluation. Many of the dipeptide sequences were derived from bioactive peptides known to undergo C-terminal processing, such as angiotensin and bradykinin, while additional sequences were incorporated to broadly sample amino acid diversity at both the P1 and P1′ positions (**Figure 1c**).

When evaluated in the absence of enzymes, several probes exhibited slow spontaneous hydrolysis under neutral aqueous conditions (**Figure S1**). This background instability depended on amino acid composition: probes bearing polar residues such as Asp or His at P1 or P1′, or Ser or Thr at P1′, were moderately unstable. However, their decomposition rates were substantially slower than those of probes containing Ser or Thr at P1, and enzyme-dependent activity could be reliably quantified by subtracting the background fluorescence increase during kinetic analysis.

Using this probe library, we examined the activities of representative carboxypeptidases, including prostate-specific membrane antigen (PSMA), carboxypeptidase A1 (CPA1), CPA4, carboxypeptidase B1 (CPB1), carboxypeptidase E (CPE), and carboxypeptidase M (CPM). Carboxylesterase 2 (CES2) was included as a negative control. Through this systematic profiling, we identified key sequence requirements and tolerances that govern substrate recognition by each enzyme (**Figure 1c**).

PSMA is widely recognized as a highly specific exopeptidase, and our dataset clearly reflected this stringent substrate preference^7^. Despite extensive evaluation of multiple candidate sequences, PSMA-reactive substrates were found to be extremely limited. The Glu–Glu (EE) sequence was initially designed based on structural considerations derived from well-established PSMA ligands, such as MIP-1095, which contain a terminal glutamate and additional free carboxylate groups that contribute to high-affinity binding^8,9^. We hypothesized that introducing a glutamic acid residue at the P1 position might enhance substrate recognition by mimicking these ligand features. However, the resulting EE substrate exhibited markedly reduced reactivity toward PSMA compared with Gly–Glu (GE), which has previously been reported as the only PSMA-reactive sequence identified through liquid-phase synthesis^5^ (**Figure 1d**). In parallel, a broader set of substrate variants was synthesized on solid support and evaluated under identical assay conditions. None of these sequences (Ala–Glu, Phe–Glu, and Arg–Glu) showed detectable PSMA activity, indicating that further sequence diversification using a solid-phase approach did not yield improved substrates (**Figure 1c**). Collectively, these results **validate** the exceptionally narrow substrate tolerance of PSMA. While further sequence diversification did not yield a superior substrate to GE, this exhaustive profiling provides chemical evidence for the stringent substrate-recognition architecture of PSMA, reinforcing the necessity of precise motif selection for this specific enzyme. In contrast, other carboxypeptidases exhibited markedly broader and more tunable substrate profiles, as discussed below.

Carboxypeptidases of the A-type (CPA) comprise several isoforms, among which CPA1 and CPA2 are predominantly expressed in the pancreas and have been proposed as highly specific circulating markers for pancreatic disorders^10^. To examine their substrate preferences within the ProTide framework, we analyzed recombinant CPA1 and CPA4. Although CPA enzymes have traditionally been described as favoring aromatic residues at the P1′ position^10^, our profiling revealed that sequences such as Ala–Leu (AL) and Ala–Val (AV) were also efficiently hydrolyzed (**Figure 1c, S2**). These results indicate that CPA isoforms possess broader substrate tolerance than previously appreciated. Moreover, subtle differences in side-chain size and hydrophobicity could be exploited to distinguish the activities of individual isoforms, such as CPA1 (preferring Gly-Phe and Ala-Leu) and CPA4 (preferring Phe-Leu).

We further investigated the substrate scope of B-type carboxypeptidases, focusing on CPB1, CPE, and CPM, which play major roles in peptide hormone processing and extracellular peptide turnover^10^. As expected, these enzymes strictly recognized peptides containing basic amino acids (Arg or Lys) at the P1′ position (**Figure 1c, S3**). However, distinct preferences emerged with respect to Arg versus Lys selection and their dependence on the P1 residue. The pancreatic enzyme CPB1 showed a clear preference for Arg over Lys, whereas CPE, which is primarily expressed in the brain and neuroendocrine tissues, and CPM, which participates in the metabolism of diverse bioactive peptides at the cell surface, accepted substrates bearing either Arg or Lys at the P1′ position. In contrast, none of the three enzymes favored substrates containing polar amino acids at the P1 position, and CPE exhibited a specific preference for substrates bearing Ala at P1. These differences in substrate recognition enabled the identification of peptide sequences capable of selectively reporting the activities of individual CPB-family enzymes.

### Single-molecule carboxypeptidase activity detection and application to pancreatic cancer-associated biomarkers

Building on the substrate preferences defined above, we next sought to develop an assay capable of discriminately detecting CPs having overlapping activities, and we tried to establish this by using the single-molecule enzyme activity assay^11,12^. In single-molecule enzyme activity profiling (SEAP) platform^13^, enzyme solution is loaded into the microfabricated chamber device with concentration in that 0 or 1 molecule of the target enzyme is theoretically loaded into each femtoliter volume chamber^14^. In each chamber, the activity of individual enzyme molecule is analyzed using multi-colored fluorogenic probes (**Figure 2a**). Besides the high detection sensitivity, the assay platform performed with multi-colored substrates can discriminate resembling enzyme subtypes and proteoforms based on the different reactivities toward multiple substrates^13^. The feature is advantageous to characterize and monitor the disease-related alterations of specific enzyme activities in blood samples for disease diagnosis. In particular, when the enzyme of interest is expressed in a tissue-restricted manner, its circulating activity can serve as a direct molecular indicator of the tissue-of-origin. Given that several CPA/CPB isoforms are predominantly expressed in the pancreas^10^, we hypothesized that detecting their activity at the single-molecule level could enable sensitive assessment of pancreatic pathophysiology.

The microdevice-based single-molecule assay imposes stringent requirements on fluorogenic probes, including high brightness, appropriate excitation/emission wavelengths, and sufficient hydrophilicity^13,15,16^. The commonly used fluorophore 4-methylumbelliferone (4MU) did not meet these criteria and was therefore unsuitable for this platform^13^.

**Figure 2.**
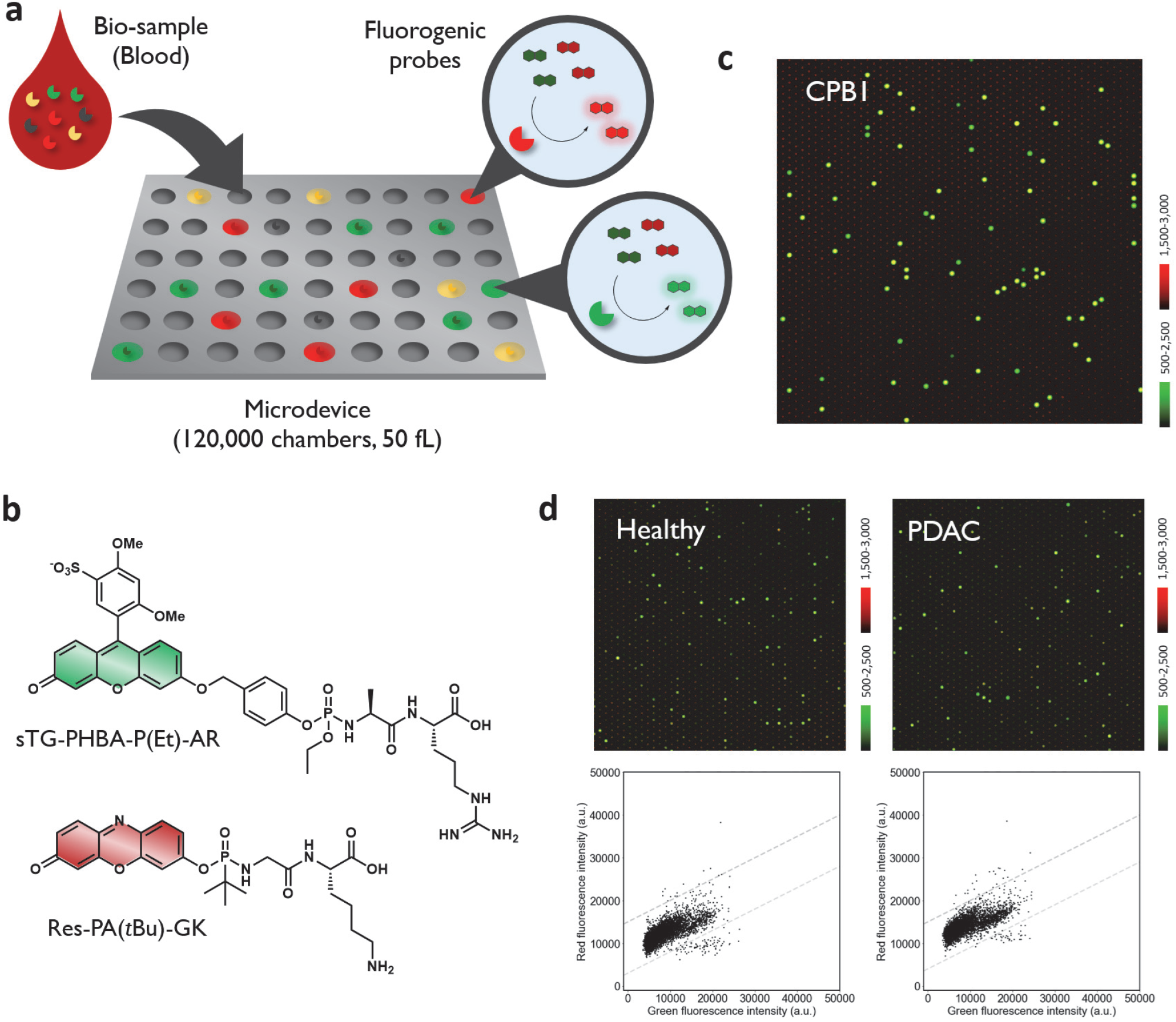
Multi-color single-molecule carboxypeptidase assay to acquire the pathological activity patterns of enzymes. (a) Concept of SEAP platform for analyzing single-molecule carboxypeptidase activities at proteoform resolution. (b) Structures of sTG-PHBA-P(Et)-AR and Res-PA(*t*Bu)-GK. (c) Fluorescence images of microdevice after loading sTG-PHBA-P(Et)-AR (green, 10 μM) and Res-PA(*t*Bu)-GK (red, 30 μM) with trypsin-activated recombinant CPB1 (0.1 ng/mL) in HEPES-Na buffer (100 mM, pH 7.4) containing ZnCl_2_ (10 μM), CaCl_2_ (1 mM), MgCl_2_ (1 mM), DTT (100 μM), Triton X-100 (150 μM), and IRDye800 (30 μM) and incubated at 25°C for 24 h. (d) Fluorescence images of microdevice after loading sTG-PHBA-P(Et)-AR (10 μM) and Res-PA(*t*Bu)-GK (30 μM) with blood samples (1/20,000) and trypsin (recombinant, 1.66 μg/mL) in HEPES-Na buffer (100 mM, pH 7.4) containing ZnCl_2_ (10 μM), CaCl_2_ (1 mM), MgCl_2_ (1 mM), DTT (100 μM), Triton X-100 (150 μM), and IR-Dye 800 (30 μM) and incubated at 25°C for 24 h. Dot plot indicates the distribution of the activity species, each dot corresponding to single-molecule enzyme spots with green fluorescence (x-axis) and red fluorescence (y-axis).

To address these limitations, we designed green-and red-emitting fluorogenic probes using sulfonated TokyoGreen (sTG)^13^ and resorufin (Res), respectively. However, the phenolic groups of these fluorophores are more acidic than that of 4MU, rendering ProTide probes with direct fluorophore attachment chemically unstable. To improve probe stability while retaining efficient signal generation, we implemented two complementary design strategies. In design A, the fluorophore was connected to the peptide substrate via a self-immolative *p*-hydroxybenzyl alcohol (PHBA) linker, allowing fluorophore release upon enzymatic cleavage while minimizing premature hydrolysis. In design B, we replaced the conventional ethyl phosphoroamidate-based ProTide with a more stable *tert*-butyl phosphonoamidate-based ProTide scaffold^5^. Both strategies were compatible with sTG and resorufin fluorophores and enabled robust probe synthesis by solid-phase methods (**Scheme S3**-**S5**). Based on the structure–activity relationships established in bulk enzymatic assays, we selected probe pairs targeting CPB (sTG–PHBA–P(Et)-GR and Res–PA(*t*Bu)-AK) and CPA (sTG– PHBA–P(Et)-AY and Res–PHBA–P(Et)-GL) for subsequent single-molecule analysis. Despite minor variations in scaffold architecture, all probes were designed according to the same underlying principles of stability, fluorogenic efficiency, and enzyme selectivity required for microdevice-based measurements (**Figure 2b**). As expected, the probe sets were able to discriminate between different carboxypeptidases (CPB1, CPE, CPM, CPA1, CPA2, and CPA4) in SEAP platforms (**Figure 2c, S4, S5**). Finally, we have analyzed plasma samples of human subjects to see if we can discriminately detect physiological CP activities in blood. In order to detect various carboxypeptidase species including preproenzymes^10^, we have activated them by treating blood samples with trypsin. CPB-selective probes predominantly reported activity attributable to liver-derived CPB family members, with only a minor contribution from pancreatic CPB1, indicating that CPB activity in circulation is largely non-specific with respect to pancreatic disease (**Figure 2d**). In contrast, CPA-selective probes revealed a well-resolved population of enzyme molecules (**Figure 3a, 3b**), and the levels of clusters corresponding to pancreas-specific carboxypeptidases, CPA1 (cluster #5) and CPA2 (cluster #3 and #4), were significantly elevated in a subset of samples from individuals with pancreatic cancer. The result was in a sharp contrast with the ubiquitous enzyme CPA4 (cluster #2), in which the elevation was observed both in healthy controls and pancreatic cancer patients (**Figure 3c**).

**Figure 3.**
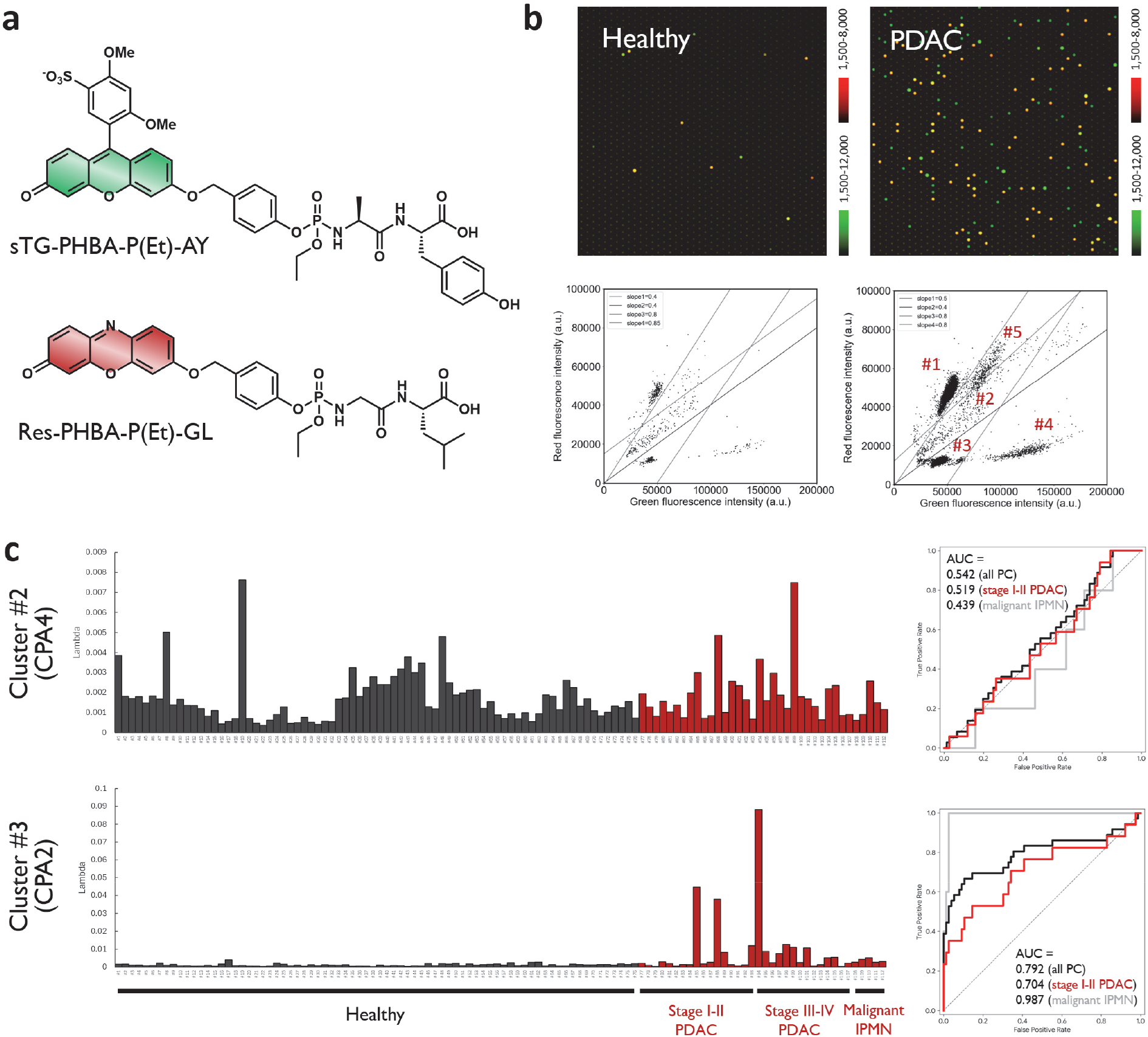
Pancreas-specific CPA subtypes serve as the potential biomarker of pancreatic cancer. (a) Structures of sTG-PHBA-P(Et)-AY and Res-PHBA-P(Et)-GL. (c) Fluorescence images of microdevice after loading sTG-PHBA-P(Et)-AY (green, 10 μM) and Res-PHBA-P(Et)-GL (red, 10 μM) with blood samples (1/2,500) and trypsin (1.66 μg/mL) in Tris-HCl buffer (100 mM, pH 8.5) containing ZnCl_2_ (10 μM), CaCl_2_ (1 mM), MgCl_2_ (1 mM), DTT (100 μM), Triton X-100 (150 μM), and IR-Dye 800 (30 μM) and incubated at 25°C for 24 h. Dot plot indicates the distribution of the activity species, each dot corresponding to single-molecule enzyme spots with green fluorescence (x-axis) and red fluorescence (y-axis). (c) Lambda values (possibility of having one enzyme in the chamber) of clusters corresponding to ubiquitously expressed CPA4 (cluster #2 of (b)) and pancreas-specific CPA2 (cluster #3 of (b)). ROC curves were constructed based on the lambda values. Black line indicates the results of all pancreatic cancer (PC) patients, Red line indicates the results of stage I-II pancreatic adenocarcinoma (PDAC), and gray line indicates the results of malignant IPMN (IPMN-HGD, IPMN-INV). Whole data is shown in **Figure S6-S10.** Independent ROC curves are shown in **Figure S11**.

We conducted an analysis of plasma samples from 112 individuals, including 76 healthy controls and 36 pancreatic cancer patients (31 pancreatic ductal adenocarcinoma (PDAC) and 5 malignant intraductal papillary mucinous neoplasm (IPMN), including high-grade dysplasia (IPMN-HGD) and invasive carcinoma (IPMN-INV); see **Table S2, S3**). Our analysis revealed that the population assigned as CPA2 with low activity (cluster #3 in **Figure 3b**) provided the best discrimination between pancreatic cancer patients and healthy subjects, with an AUC value of 0.792 (**Figure 3c, S6**-**S10**). This molecular species exhibited a substrate preference comparable to recombinant CPA2 but showed weaker catalytic activity (**Figure S5**), indicating that these species are likely unique proteoforms of CPA2 with post-translational modifications that lower their activity. Notably, high diagnostic performance was maintained even when the analysis was restricted to stage I–II PDAC (n = 20, AUC = 0.704), suggesting that pancreas-derived CPA activities can be detected at early disease stages. Furthermore, for malignant IPMN (comprising IPMN-HGD and IPMN-INV), we observed a remarkably high AUC of 0.987. Although this result is based on a limited cohort (n = 5), it underscores the potential of our platform for the early detection of malignant IPMN, which is critical for timely surgical resection. These results highlight the potential of pancreas-derived CPAs as minimally invasive biomarkers for the early detection of pancreatic cancer, complementing existing diagnostic workflows that currently lack sensitivity in the early disease window^17^.

## Discussion

Our study identifies circulating carboxypeptidase activities as a previously unrecognized class of activity-based biomarkers for pancreatic cancer and establishes a generalizable synthetic and analytical framework for their detection at the single-molecule level. Enzymes with tissue-restricted expression profiles represent compelling biomarker candidates, as alterations in their circulating activity may directly reflect organ-specific pathological processes. In this context, several carboxypeptidases are predominantly expressed in the pancreas, positioning them as a promising yet previously underexplored source of activity-based biomarkers^3^. The ability to detect their activity sensitively in blood thus enables a diagnostic strategy with intrinsic specificity for pancreatic function and dysfunction.

Beyond the direct diagnostic implications, this work presents a systematic structure–activity relationship (SAR) map of substrate preferences across a broad panel of carboxypeptidase family members, encompassing both metallo-and serine-type enzymes. By leveraging a solid-phase ProTide synthesis platform, we rapidly generated and evaluated a diverse library of peptide-based probes, enabling the identification of distinct and, in some cases, noncanonical substrate specificities. These findings provide a conceptual and practical foundation for future development of carboxypeptidase-targeting molecular tools, including fluorogenic and luminescent probes, selective imaging agents, and prodrug designs. The SAR knowledge base established here is expected to facilitate the rational selection or engineering of substrates capable of discriminating among closely related isoforms in complex biological environments. Intriguingly, our single-molecule analysis revealed that the most effective discrimination of pancreatic cancer was achieved by a specific CPA2 sub-population (cluster #3) exhibiting lower catalytic activity than the recombinant enzyme. The detection of such a well-resolved, low-activity population highlights the unique capability of the SEAP platform to distinguish specific enzyme proteoforms that might otherwise be masked in bulk assays^11^. While the exact molecular nature of this species remains to be determined, its prevalence in pancreatic cancer patients suggests that disease-specific post-translational modifications or altered activation dynamics may generate unique enzymatic signatures, a phenomenon previously observed in other biomarker enzyme systems^13,18,19^.

A current limitation of this study is that the mechanistic basis for altered circulating carboxypeptidase activity in pancreatic cancer remains unresolved. Prior proteomic analyses have reported aberrant C-terminal processing of apolipoprotein A-II (ApoA-II) in pancreatic cancer^20–23^, suggesting dysregulated carboxypeptidase function. While our findings are consistent with these observations, the causal relationships among pancreatic tissue alterations, enzyme expression or activation dynamics, and the generation of specific proteolytic signatures remain to be elucidated. Clarifying these mechanisms through biochemical and histopathological studies will be essential for determining the temporal and cellular origins of the observed enzymatic changes, particularly in the context of early disease development.

Taken together, our results expand the repertoire of enzymes amenable to single-molecule activity-based detection and highlight the diagnostic and mechanistic potential of carboxypeptidases in pancreatic cancer. By integrating scalable probe synthesis, global enzymatic profiling, and ultra-sensitive activity readouts, this work establishes a platform for discovering biomarker enzymes whose activities are tightly linked to organ-specific pathophysiology.

## Conclusion

In summary, we developed a solid-phase ProTide probe synthesis strategy that enables systematic exploration of carboxypeptidase substrate selectivity and facilitates the creation of single-molecule enzyme activity assays. Using this platform, we identified circulating CPA activity as a sensitive biomarker with strong pancreatic relevance, capable of distinguishing pancreatic cancer patients from healthy individuals, even at early disease stages. These findings not only advance the molecular understanding of carboxypeptidase biology but also provide a foundation for next-generation diagnostic approaches that directly quantify enzyme activity as a readout of disease-associated tissue states.

## AUTHOR INFORMATION

### Competing Financial Interests

Y. K. is a cofounder, employee and shareholder of Cosomil, Inc. M. M. and H. H.. are employees of Cosomil, Inc. T. K. and K. H. are advisors and shareholders of Cosomil, Inc. T. K. and Y. U. are inventors of a patent describing the design strategies of fluorogenic probes. This study was supported in part by Cosomil, Inc. through a collaborative research agreement with the University of Tokyo.

### Author Contributions

T. K. and Y. U. conceived the experimental design. T. K., and T. U. constructed solid-phase-based synthetic schemes. T. K., T. U., M. H., and S. K. synthesized and characterized 4MU-based probes. T. K. and M. M. synthesized and characterized probes for single-molecule analysis. T. K., M. H., and S. K. acquired experimental data of 4MU-based probes. M. M., and H. H. acquired experimental data using blood samples. The experimental data were analyzed under the supervision of T. K., Y. K., K. H., and Y. U. The manuscript was written by T. K. All authors have given approval to the final version of the manuscript.

## Acknowledgements

We thank Dr. Yugo Kuriki, Dr. Shingo Sakamoto, and Dr. Kazuki Takahashi for fruitful discussions about the project. This work was financially supported by MEXT (20H04694, 21A303, 22H02217, 23K23484, 25K01911 and 25K22520 to T. K.), JST (PRESTO (13414915), PRESTO Network (17949814) and FOREST (24012649) to T. K., START (20353017) to T. K., and K. H.), and AMED (FORCE (22581634) and P-PROMOTE (25131640) to T. K. and K. H., Research on Development of New Drugs (23809006) for T. K., P-PROMOTE (18cm0106403h0003) to K. H., P-CREATE (25ama221431h0002) to K. H. T. K. received support from the Naito Foundation, The Mochida Memorial Foundation for Medical and Pharmaceutical Research, Chugai Foundation for Innovative Drug Discovery Science, MSD Life Science Foundation, Hoansha Foundation, and University of Tokyo Gap Fund Program.

