## Supplementary Information for "Solid-Phase Synthesis of ProTide Fluorogenic Probes Enables Systematic Profiling of Carboxypeptidase Activity"

**Contents**

**Methods**
**Supplementary data**
**Supplementary data for synthesis and characterization of compounds**
**Supplementary references**

### Methods

#### Materials

| REAGENT OR RESOURCE | SOURCE | IDENTIFIER |
|---|---|---|
| PSMA/FOLH1 (*homo sapiens*) | R&D Biosystems | Cat. #4234-ZN<br>Lot #QBU1624011 |
| Carboxypeptidase A1 (CPA1, *homo sapiens*) | R&D Biosystems | Cat. #2856-ZN<br>Lot #OAK0421021 |
| Carboxypeptidase A2 (CPA2, *homo sapiens*) | R&D Biosystems | Cat. #2896-ZN<br>Lot #OFM0225051 |
| Carboxypeptidase A4 (CPA4, *homo sapiens*) | R&D Biosystems | Cat. #5906-ZN<br>Lot #TIV0423091 |
| Carboxypeptidase B1 (CPB1, *homo sapiens*) | R&D Biosystems | Cat. #2897-ZN<br>Lot #OKA0224031 |
| Carboxypeptidase E (CPE, *homo sapiens*) | R&D Biosystems | Cat. #3587-ZN<br>Lot # PEK0123081 |
| Carboxypepidase M (CPM, *homo sapiens*) | R&D Biosystems | Cat. #7457-ZN<br>Lot # DCEI0223121 |
| Carboxypeptidase 2 (CES2, *mus musculus*) | R&D Biosystems | Cat. #5280-CE<br>Lot #SJF0114051 |
| Trypsin, recombinant | Roche | Cat. #06369880103<br>Lot #80223100 |
| Trypsin, recombinant | Yeasen | Cat. #20416ES60<br>Lot #R6519170 |
| Trypsin, proteomics grade | Roche | Cat. #3708985001 |

**Plasma samples and ethics statement**

The blood samples used for the results reported in this article were obtained from the Okayama University Hospital Biobank (Okadai Biobank) and Japan Institute for Health Security Biobank, Japan, and all experiments using these samples were conducted at Cosomil Inc. Ethical approval for this study was obtained from the Craif Institutional Review Board (IEF-24012401). Disease stage was assigned according to the Union for International Cancer Control (UICC) TNM classification, 8th edition. During the developmental phase of the assay, plasma samples from healthy human subjects were used for the screening of the candidate probes. These samples were collected from the Kagoshima Prefectural Comprehensive Health Center through the Program for Promotion of Fundamental Studies in Health Sciences conducted by the National Institute of Biomedical Innovation of Japan, Health and Labour Sciences Research Grants from the Ministry of Health, Labour and Welfare of Japan, and P-CREATE of the Japan Agency for Medical Research and Development (AMED). Data obtained from these developmental studies are not included in the results presented in this article. Ethical approval for these studies was obtained from the central ethics committees of Nippon Medical School (M-2021-002) and the ethical committee of Nippon Medical School (A-2020-032 and A-2020-044).

**Instruments**

NMR spectra were recorded on a JEOL JNM-LA400 instrument at 400 MHz for $^{1}H$ NMR and at 100 MHz for $^{13}C$ NMR. Mass spectra (MS) were measured with a JEOL JMS-T100LC AccuToF (ESI). LC-MS analyses were performed on a Waters Acquity UPLC (H class)/QDa quadrupole MS analyzer or Acquity UPLC (H class)/Xevo TQD quadrupole MS/MS analyzer equipped with an Acquity UPLC BEH C18 column (Waters). Column chromatography using silica gel was performed on a MPLC system (Yamazen Smart Flash EPCLC AI-5805 (Tokyo, Japan)). Reversed-phase MPLC purification was performed on an Isolera One (Biotage) equipped with a SNAP Ultra C18 30 g (Biotage). UV-Visible absorbance was obtained on a Shimadzu UV-1800.

**Activation of recombinant CPB1**

Recombinant CPB1 (10 μg/mL) was mixed with trypsin (proteomics grade, Roche, 10 μg/mL) in HEPES-Na buffer (100 mM, pH 7.4) containing $CaCl_2$ (10 mM) and CHAPS (0.1%) and incubated at 25°C for 40 min. The sample was diluted by the reaction buffer and used. The concentrations described in the manuscript indicates those before the activation.

**Activation of CPs in plasma samples**

Plasma samples (1/2,500 for CPAs and 1/20,000 for CPBs) were mixed with trypsin (recombinant, from Roche for CPAs or Yeasen for CPBs, 1.66 μg/mL) in assay buffer and immediately mixed with fluorogenic probes for loading into microdevice.

**Fluorescence microscopy**

Fluorescence images were acquired by fluorescence microscope (Ti2, Nikon) equipped with a 20× dry objective lens (Plan Apo 20×), sCMOS camera (ORCA-Fusion C14440, Hamamatsu Photonics), white LED illumination unit (X-Cite Xylis, Opto Science), moving stage and rotating stage. The stage was custom-made to hold Simoa disk (Quanterix) and rotate it using motorized rotation stage(OSMS-60YAW, Opto Sigma). For continuous monitoring of multiple device images, the rotating stage rotated the disk by 15° for each acquisition. Each image was acquired in tile scan mode of 3×4 (with overlap of 1%) with perfect focus. The assay was performed using a solution containing IR-dye 800 (10 μM) as the internal standard, and the focus was adjusted using its fluorescence. All operations were controlled by NIS-Element software (Nikon). The excitation and emission filters used were FITC (mirror = 510 nm, Ex. = 460-500 nm, Em. = 510-560 nm), mCherry (mirror = 600 nm, ex. = 550-590 nm, Em. = 608-683 nm), and Cy7 (mirror = 743 nm, Ex. = 743 nm, Em. = 767 nm), respectively. Fluorescence signals of sTG-based probes were acquired with FITC filter and those of resorufin-based probes were acquired with mCherry filter. Fluorescence signals of IR-dye 800 were acquired with Cy7 filter.

**Image processing**

Images were processed using the GA3 module of NIS Elements software (Nikon). First, all fluorescence images were background-corrected using a rolling ball correction (diameter = 3 μm). Then, ROIs were chosen by bright spot detection using FITC and mCherry filters (diameter = 3 μm). The fluorescence signals were acquired as the mean of the signals from each ROI. The dot plots were generated using Python.

### Supplementary data

**Table S1.** List of known enzymes having carboxypeptidase activities. Information was extracted and summarized from the reference[1].

| Enzyme | EC | Gene | Substrate preference at C-terminal | Tissue | Physiological substrate | Optimum pH | Disease and biological phenomena |
|---|---|---|---|---|---|---|---|
| Carboxypeptidase A | 3.4.17.1 | CPA1 | Hydrophobic amino acid | Pancreas | | | Pancreatitis, Pancreatic cancer |
| Carboxypeptidase A2 | 3.4.17.15 | CPA2 | Hydrophobic amino acid | Pancreas | | | Pancreatitis, Pancreatic cancer |
| Carboxypeptidase A3 | 3.4.17.1 | CPA3 | Hydrophobic amino acid | Mast cells | | | Mast cell degranulation |
| Carboxypeptidase A4 | 3.4.17.1 | CPA4 | Hydrophobic amino acid | Various | | | |
| Carboxypeptidase A5 | 3.4.17.1 | CPA5 | Hydrophobic amino acid | Testis | | | |
| Carboxypeptidase A6 | 3.4.17.1 | CPA6 | Hydrophobic amino acid | Various | | | |
| Carboxypeptidase U | 3.4.17.20 | CPU | Arg, Lys | | | | Blood coagulation |
| Carboxypeptidase B | 3.4.17.2 | CPB1 | Arg, Lys | Pancreas | | | Pancreatitis, Pancreatic cancer |
| Lysine carboxypeptidase<br>Carboxypeptidase N | 3.4.17.3 | CPN1<br>/CPN2 | Arg, Lys | Liver | | | Breast cancer |
| Carboxypeptidase E | 3.4.17.10 | CPE | Arg, Lys | Brain | Enkephalin | | Various cancers |
| Carboxypeptidase Z | | CPZ | Arg | Female tissues | | | |
| Metallocarboxypeptidase D | 3.4.17.22 | CPD | Arg, Lys | Various | | | |
| Carboxypeptidase M | 3.4.17.12 | CPM | Arg, Lys | Various | Bradykinin,<br>kallidin,<br>enkephalin, | | Various cancers, pneumonia, cell differentiation |
| Tubulinyl-Tyr carboxypeptidase | 3.4.17.17 | VASH1<br>/VASH2 | Glu-Tyr, Glu-Phe | Various | Tubulin | | Spindle function and chromosome segregation, angiogenesis |
| Tubulin-glutamate carboxypeptidase | 3.4.17.24 | AGBL1 | Glu | Digestive tract | | | |
| Lysosomal carboxypeptidase A<br>Serine carboxypeptidase A<br>Cathepsin A | 3.4.16.5 | CTSA<br>/SCPEP1 | Hydrophobic amino acid | Various | Substrance P,<br>Enkephalin,<br>Oxytocin | Acidic | Galactosialidosis |
| Lysosomal Pro-Xaa carboxypeptidase<br>Angiotensinase C | 3.4.16.2 | PRCR | Pro-Xaa | Various | Angiotensin II | Acidic | |
| Pepdiyl-dipeptidase A<br>Angiotensin I-converting enzyme | 3.4.15.1 | ACE | C-term dipeptide | Various | Angiotensin | | High blood pressure |
| Angiotensin-converting enzyme 2 | 3.4.17.23 | ACE2 | C-term dipeptide | Various | Angiotensin | | |
| Glutamate carboxypeptidase II<br>Prostate specific membrane antigen | 3.4.17.21 | FOLH1 | Glu | Various | NAAG, Folic acid | | Prostate cancer |

**Table S2.** List of blood samples of healthy subjects. Source, and measured lambda values of cluster #1-#5 are shown. OBB indicates the blood samples from Okayama University Hospital Biobank (Okadai Biobank) and NCGM indicates the blood samples from Japan Institute for Health Security Biobank.

| Label | Stage | Source | Cluster #1 | Cluster #2 | Cluster #3 | Cluster #4 | Cluster #5 |
|---|---|---|---|---|---|---|---|
| Control | | NCGM | 0.005883 | 0.003855 | 0.001592 | 0.000444 | 0.000411 |
| Control | | NCGM | 0.004148 | 0.00181 | 0.001686 | 0.00091 | 0.000112 |
| Control | | NCGM | 0.001608 | 0.001709 | 0.001173 | 0.000201 | 0.000101 |
| Control | | NCGM | 0.00242 | 0.001797 | 0.00122 | 0.000884 | 0.000121 |
| Control | | NCGM | 0.002101 | 0.001482 | 0.000913 | 0.000435 | 0.000268 |
| Control | | NCGM | 0.002233 | 0.001836 | 0.001022 | 0.000416 | 0.000312 |
| Control | | NCGM | 0.004268 | 0.001675 | 0.002342 | 0.000657 | 0.000317 |
| Control | | NCGM | 0.002122 | 0.005021 | 0.001092 | 0.000323 | 0.000428 |
| Control | | NCGM | 0.004512 | 0.001395 | 0.001524 | 0.000336 | 0.000257 |
| Control | | NCGM | 0.001557 | 0.001661 | 0.000836 | 0.000564 | 0.000397 |
| Control | | NCGM | 0.001847 | 0.001359 | 0.00088 | 0.000436 | 0.000322 |
| Control | | NCGM | 0.000905 | 0.001353 | 0.000629 | 0.000257 | 8.58E-05 |
| Control | | NCGM | 0.001211 | 0.001202 | 0.000656 | 0.000219 | 0.000227 |
| Control | | NCGM | 0.002107 | 0.00082 | 0.001093 | 0.000364 | 0.000148 |
| Control | | NCGM | 0.002033 | 0.001104 | 0.000535 | 0.00036 | 0.000184 |
| Control | | NCGM | 0.003785 | 0.000435 | 0.001725 | 0.000469 | 0.000184 |
| Control | | NCGM | 0.006771 | 0.000554 | 0.004116 | 0.000538 | 0.000311 |
| Control | | NCGM | 0.002254 | 0.000704 | 0.000964 | 0.000369 | 0.000218 |
| Control | | NCGM | 0.001651 | 0.007629 | 0.001031 | 0.000863 | 0.000738 |
| Control | | NCGM | 0.001097 | 0.000712 | 0.00067 | 0.000243 | 0.000159 |
| Control | | NCGM | 0.001581 | 0.000462 | 0.000613 | 0.000409 | 0.000129 |
| Control | | NCGM | 0.001542 | 0.00036 | 0.000318 | 0.000268 | 9.22E-05 |
| Control | | NCGM | 0.002452 | 0.000628 | 0.000705 | 0.000301 | 0.00012 |
| Control | | NCGM | 0.004524 | 0.000912 | 0.002348 | 0.0008 | 0.000215 |
| Control | | NCGM | 0.002863 | 0.001283 | 0.001587 | 0.000743 | 0.000236 |
| Control | | NCGM | 0.001416 | 0.000461 | 0.000787 | 0.00041 | 0.000142 |
| Control | | NCGM | 0.002324 | 0.000502 | 0.001162 | 0.000418 | 0.000117 |
| Control | | NCGM | 0.001537 | 0.000765 | 0.000554 | 0.000244 | 9.24E-05 |
| Control | | NCGM | 0.003074 | 0.000519 | 0.001323 | 0.000645 | 0.000126 |
| Control | | NCGM | 0.0031 | 0.000411 | 0.000964 | 0.000318 | 0.000159 |
| Control | | NCGM | 0.003683 | 0.000569 | 0.001381 | 0.000594 | 0.000201 |
| Control | | NCGM | 0.002338 | 0.000545 | 0.001139 | 0.000402 | 0.000101 |
| Control | | NCGM | 0.002366 | 0.001674 | 0.001095 | 0.000393 | 0.000114 |
| Control | | NCGM | 0.003501 | 0.001738 | 0.001327 | 0.000787 | 0.000146 |
| Control | | NCGM | 0.002981 | 0.003244 | 0.00057 | 0.000649 | 0.000281 |
| Control | | NCGM | 0.00165 | 0.001801 | 0.000645 | 0.00031 | 0.000134 |
| Control | | NCGM | 0.00124 | 0.002254 | 0.000628 | 0.000478 | 0.000142 |
| Control | | NCGM | 0.00123 | 0.002795 | 0.000243 | 0.000234 | 0.000117 |
| Control | | NCGM | 0.003173 | 0.002403 | 0.001214 | 0.000419 | 0.000293 |
| Control | | NCGM | 0.002006 | 0.002403 | 0.000762 | 0.000212 | 0.000186 |
| Control | | NCGM | 0.002506 | 0.002808 | 0.000981 | 0.000687 | 0.000126 |
| Control | | NCGM | 0.003216 | 0.003174 | 0.000971 | 0.000301 | 0.000218 |
| Control | | NCGM | 0.002186 | 0.003786 | 0.000394 | 0.000369 | 0.000226 |
| Control | | NCGM | 0.002237 | 0.002999 | 0.000829 | 0.000335 | 0.000235 |
| Control | | NCGM | 0.001787 | 0.003473 | 0.000772 | 0.000696 | 0.000117 |
| Control | | NCGM | 0.004231 | 0.001299 | 0.002053 | 0.000419 | 0.000193 |
| Control | | NCGM | 0.001704 | 0.001189 | 0.00086 | 0.000169 | 5.90E-05 |
| Control | | NCGM | 0.003525 | 0.004801 | 0.001999 | 0.000455 | 0.00025 |
| Control | | NCGM | 0.002712 | 0.002497 | 0.001687 | 0.000413 | 0.000207 |
| Control | | NCGM | 0.004848 | 0.001884 | 0.002227 | 0.000745 | 0.00031 |
| Control | | NCGM | 0.002067 | 0.001985 | 0.000942 | 0.00032 | 0.000119 |
| Control | | NCGM | 0.003629 | 0.002121 | 0.001802 | 0.000394 | 0.000134 |
| Control | | NCGM | 0.002747 | 0.002153 | 0.001114 | 0.000184 | 0.000201 |
| Control | | NCGM | 0.003529 | 0.001207 | 0.001392 | 0.000486 | 0.000159 |
| Control | | NCGM | 0.002325 | 0.001553 | 0.001527 | 0.000475 | 0.00011 |
| Control | | NCGM | 0.004072 | 0.000888 | 0.001784 | 0.000804 | 0.00031 |
| Control | | NCGM | 0.002714 | 0.00072 | 0.001214 | 0.000369 | 0.00031 |
| Control | | NCGM | 0.005063 | 0.000954 | 0.002507 | 0.000819 | 0.000367 |
| Control | | NCGM | 0.001977 | 0.000695 | 0.000922 | 0.000377 | 0.00031 |
| Control | | NCGM | 0.002033 | 0.000987 | 0.001171 | 0.000686 | 0.000125 |
| Control | | NCGM | 0.007244 | 0.000552 | 0.002719 | 0.000962 | 0.000351 |
| Control | | NCGM | 0.004584 | 0.000951 | 0.002837 | 0.000285 | 0.00154 |
| Control | | NCGM | 0.001704 | 0.001889 | 0.000768 | 0.000494 | 0.000168 |
| Control | | NCGM | 0.003655 | 0.001815 | 0.001405 | 0.000778 | 0.000176 |
| Control | | NCGM | 0.001826 | 0.001148 | 0.000762 | 0.000276 | 0.000109 |
| Control | | NCGM | 0.001357 | 0.002621 | 0.000754 | 0.000544 | 0.00041 |
| Control | | NCGM | 0.003494 | 0.002251 | 0.001697 | 0.000806 | 0.000176 |
| Control | | NCGM | 0.004317 | 0.001321 | 0.001498 | 0.000732 | 0.000143 |
| Control | | NCGM | 0.002712 | 0.001698 | 0.001251 | 0.000475 | 0.000137 |
| Control | | NCGM | 0.002622 | 0.001093 | 0.001311 | 0.000286 | 9.24E-05 |
| Control | | NCGM | 0.004447 | 0.000846 | 0.001491 | 0.000653 | 0.000159 |
| Control | | NCGM | 0.003697 | 0.001014 | 0.001467 | 0.000386 | 0.000143 |
| Control | | NCGM | 0.002411 | 0.001326 | 0.001201 | 0.00113 | 0.000107 |
| Control | | NCGM | 0.003888 | 0.00124 | 0.001877 | 0.000385 | 0.000168 |
| Control | | NCGM | 0.002512 | 0.001554 | 0.001453 | 0.000731 | 0.000286 |
| Control | | NCGM | 0.003779 | 0.00073 | 0.001946 | 0.000999 | 0.000252 |

**Table S3.** List of blood samples of pancreatic cancer patients. Source, and measured lambda values of cluster #1-#5 are shown. OBB indicates the blood samples from Okayama University Hospital Biobank (Okadai Biobank) and NCGM indicates the blood samples from Japan Institute for Health Security Biobank. PC = pancreatic cancer, IPMN-HGD = intraductal papillary mucinous neoplasm (IPMN) with high-grade dysplasia, IPMN-INV = invasive carcinoma associated with IPMN.

| Label | Stage | Source | Cluster #1 | Cluster #2 | Cluster #3 | Cluster #4 | Cluster #5 |
|---|---|---|---|---|---|---|---|
| PC | IA | NCGM | 0.003236 | 0.001945 | 0.002171 | 0.001165 | 0.000126 |
| PC | IA | NCGM | 0.003812 | 0.001279 | 0.001094 | 0.000589 | 0.000311 |
| PC | IA | OBB | 0.000303 | 0.000816 | 0.000336 | 0.000193 | 0.000193 |
| PC | IA | OBB | 0.003158 | 0.001562 | 0.002926 | 0.000661 | 0.000292 |
| PC | IA | OBB | 0.003192 | 0.001031 | 0.002019 | 0.000452 | 0.000176 |
| PC | IA | OBB | 0.001273 | 0.00139 | 0.000578 | 0.000226 | 0.000117 |
| PC | IB | OBB | 0.002299 | 0.000746 | 0.001483 | 0.000434 | 8.67E-05 |
| PC | IB | NCGM | 0.004357 | 0.001986 | 0.001441 | 0.000285 | 0.000804 |
| PC | IB | NCGM | 0.055156 | 0.002998 | 0.044718 | 0.011093 | 0.008566 |
| PC | IIA | NCGM | 0.002266 | 0.000723 | 0.001839 | 0.000497 | 0.000227 |
| PC | IIA | OBB | 0.004379 | 0.000853 | 0.002729 | 0.000563 | 0.000253 |
| PC | IIA | OBB | 0.04681 | 0.004859 | 0.037879 | 0.004374 | 0.003821 |
| PC | IIB | OBB | 0.010821 | 0.001004 | 0.00826 | 0.001222 | 0.000628 |
| PC | IIB | OBB | 0.00321 | 0.002565 | 0.0014 | 0.000285 | 0.000201 |
| PC | IIB | OBB | 0.002144 | 0.002345 | 0.00072 | 0.001382 | 0.000193 |
| PC | IIB | OBB | 0.001733 | 0.001641 | 0.001297 | 0.000419 | 0.000159 |
| PC | IIB | OBB | 0.012291 | 0.000676 | 0.012028 | 0.001202 | 0.000588 |
| PC | III | NCGM | 0.092446 | 0.003666 | 0.088254 | 0.009755 | 0.019536 |
| PC | III | NCGM | 0.0136 | 0.001593 | 0.008881 | 0.001725 | 0.001188 |
| PC | III | OBB | 0.012554 | 0.002975 | 0.002489 | 0.000637 | 0.001291 |
| PC | III | OBB | 0.017037 | 0.001878 | 0.00763 | 0.000889 | 0.000696 |
| PC | III | OBB | 0.019205 | 0.001206 | 0.012751 | 0.001742 | 0.001206 |
| PC | IV | NCGM | 0.03455 | 0.007498 | 0.011002 | 0.003265 | 0.001722 |
| PC | IV | NCGM | 0.004896 | 0.001801 | 0.002624 | 0.00036 | 0.000394 |
| PC | IV | NCGM | 0.014516 | 0.001685 | 0.010728 | 0.001257 | 0.000729 |
| PC | IV | NCGM | 0.002368 | 0.001312 | 0.000665 | 0.000307 | 0.000222 |
| PC | IV | OBB | 0.003237 | 0.000647 | 0.001387 | 0.000446 | 0.000135 |
| PC | IV | OBB | 0.013633 | 0.002195 | 0.005254 | 0.000871 | 0.000788 |
| PC | IV | OBB | 0.009602 | 0.002354 | 0.005492 | 0.000904 | 0.000418 |
| PC | IV | NCGM | 0.000545 | 0.000855 | 0.00031 | 0.000151 | 0.000143 |
| PC | IV | OBB | 0.004312 | 0.000911 | 0.002299 | 0.000392 | 0.000212 |
| IPMN-HGD | 0 | OBB | 0.005814 | 0.000621 | 0.002819 | 0.000914 | 0.000285 |
| IPMN-HGD | 0 | OBB | 0.006819 | 0.000913 | 0.004306 | 0.000787 | 0.000243 |
| IPMN-INV | IA | NCGM | 0.00809 | 0.002576 | 0.005262 | 0.001074 | 0.00042 |
| IPMN-INV | IA | OBB | 0.005214 | 0.001509 | 0.002724 | 0.000847 | 0.000587 |
| IPMN-INV | IIB | OBB | 0.006176 | 0.001157 | 0.003126 | 0.000243 | 0.000226 |

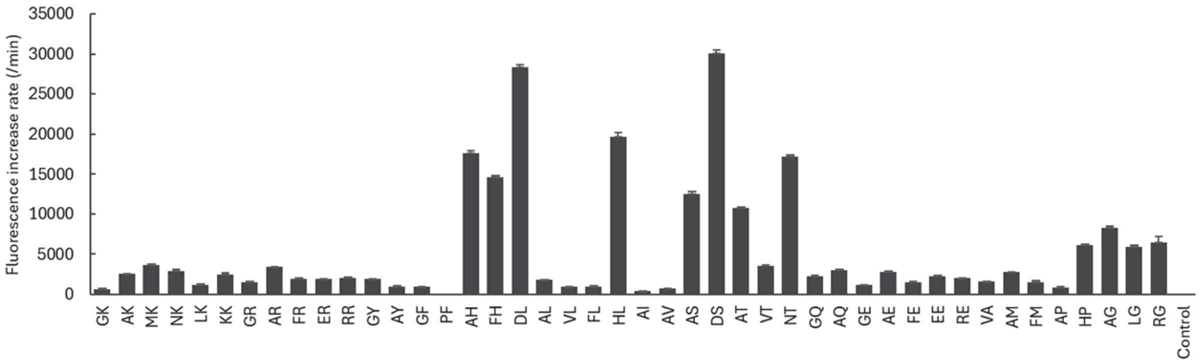


**Figure S1.** Hydrolysis of 4MU-based ProTide probes in neutral buffer. Error bars represent S. D. (n = 4). The fluorescence increase rate of 30 μM solution in Dulbecco's phosphate buffered saline (DPBS, pH 7.4) were monitored, and the slope was calculated for the data of first 60 min.

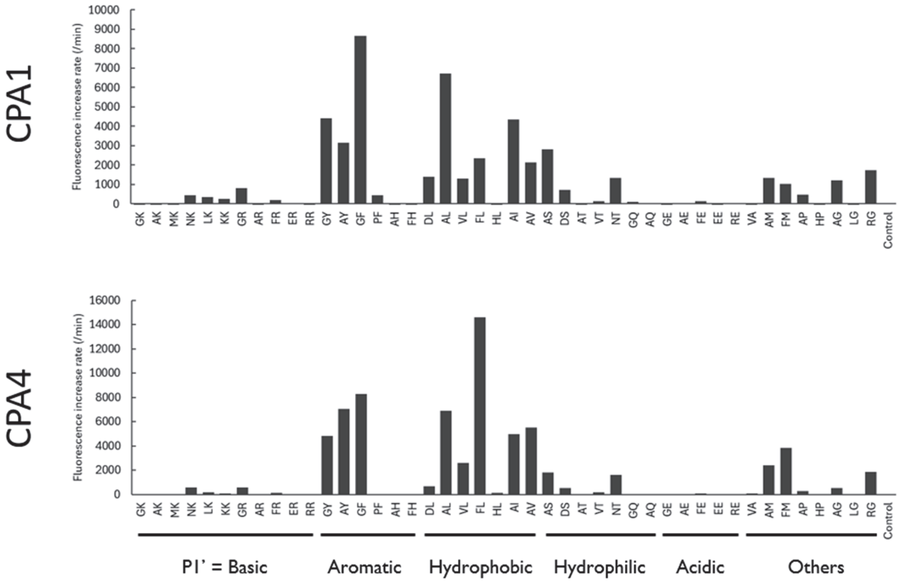


**Figure S2.** Reactivity of carboxypeptidase A. The data was derived from **Figure 1c**.

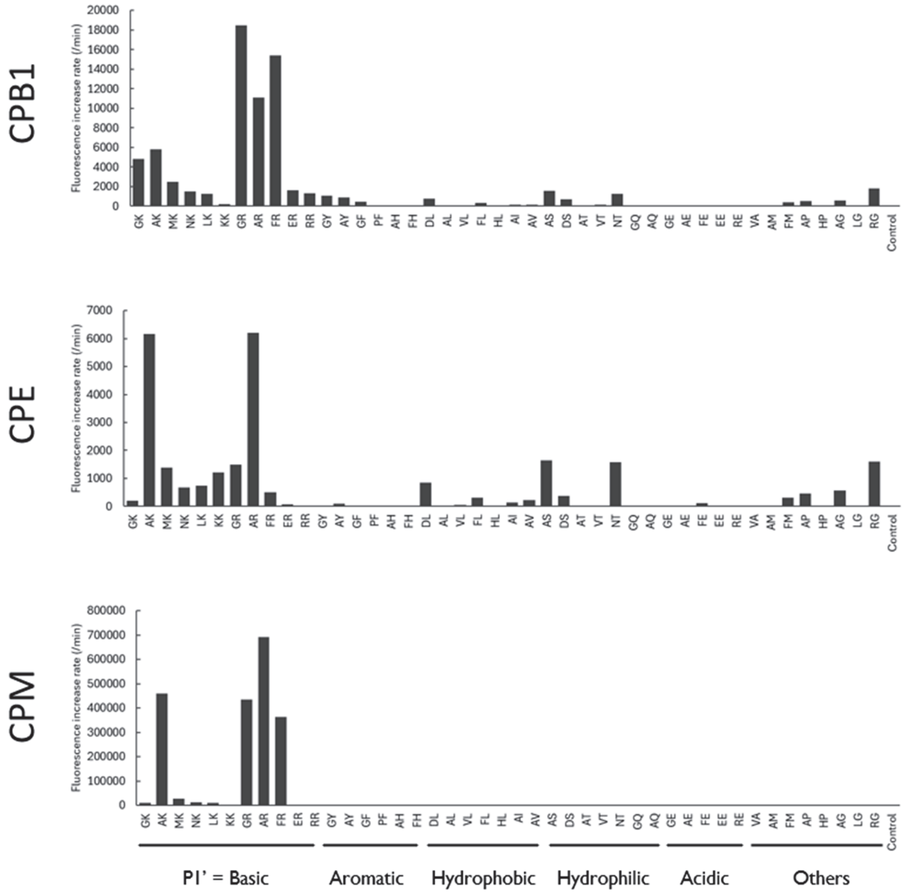


**Figure S3.** Reactivity of carboxypeptidase B. The data was derived from **Figure 1c**.

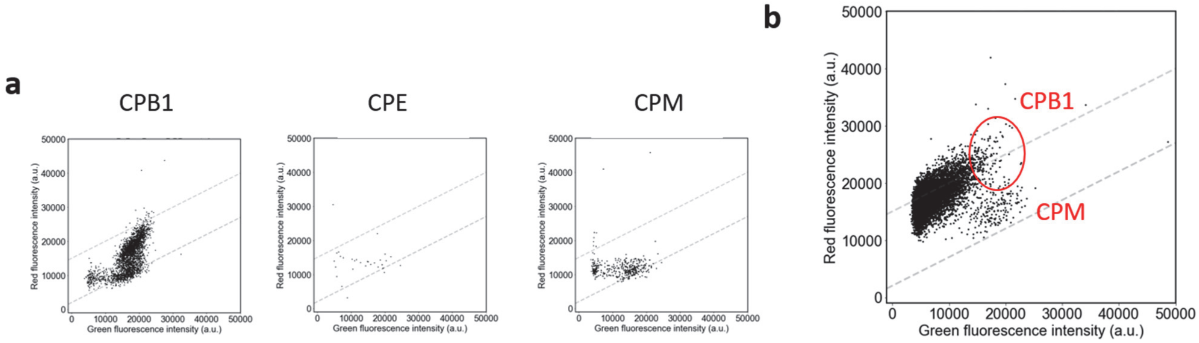


**Figure S4.** Single-molecule analysis of carboxypeptidase B. (a) Fluorescence images of microdevice were taken after loading sTG-PHBA-P(Et)-AR (green, 10 μM) and Res-PA(*t*Bu)-GK (red, 30 μM) with trypsin-activated CPB1 (0.1 ng/mL), CPE (0.1 ng/mL), or CPM (0.1 ng/mL) in HEPES-Na buffer (100 mM, pH 7.4) containing $ZnCl_2$ (10 μM), $CaCl_2$ (1 mM), $MgCl_2$ (1 mM), DTT (100 μM), Triton X-100 (150 μM), and IR-Dye 800 (30 μM) and incubating at 25°C for 24 h. (b) The data was acquired in same condition as in (a), using 1/25,000-diluted human plasma and trypsin (1.66 μg/mL).

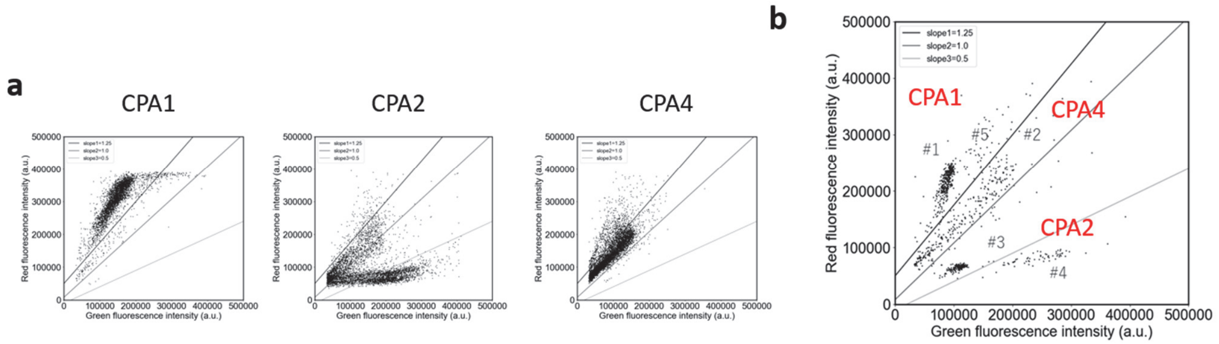


**Figure S5.** Single-molecule analysis of carboxypeptidase A. (a) Fluorescence images of microdevice were taken after loading sTG-PHBA-P(Et)-AY (green, 10 μM) and Res-PHBA-P(Et)-GL (red, 10 μM) with CPA1 (0.1 ng/mL), CPA2 (0.1 ng/mL), or CPA4 (0.1 ng/mL) in Tris-HCl buffer (100 mM, pH 8.5) containing $ZnCl_2$ (10 μM), $CaCl_2$ (1 mM), $MgCl_2$ (1 mM), DTT (100 μM), Triton X-100 (150 μM), and IR-Dye 800 (30 μM) and incubating at 25°C for 24 h. (b) The data was acquired in same condition as in (a), using 1/2,500-diluted human plasma and trypsin (1.66 μg/mL).

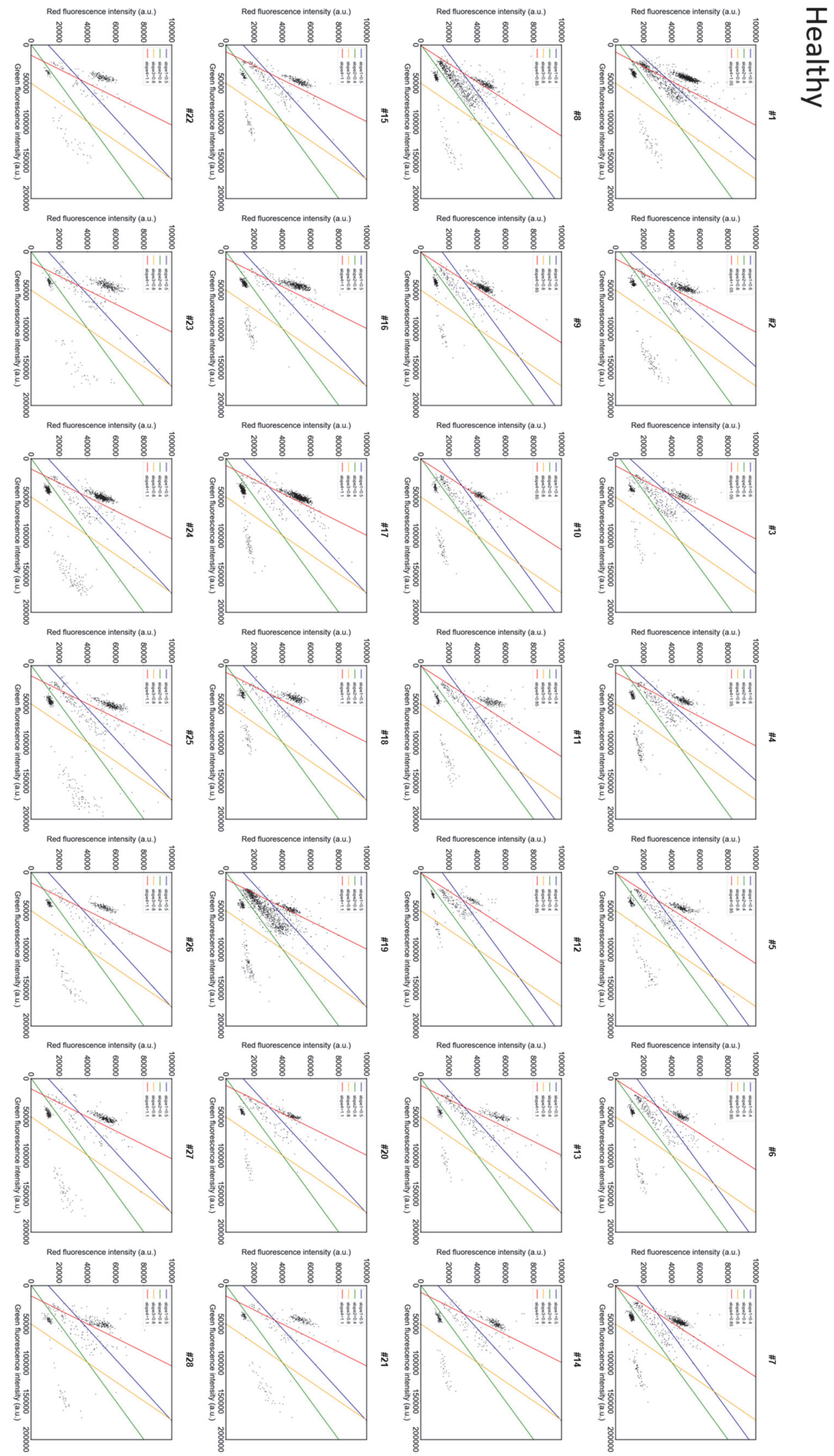


**Figure S6.** Scattered plot of single-molecule CPA activities of healthy subjects (#1-#28)

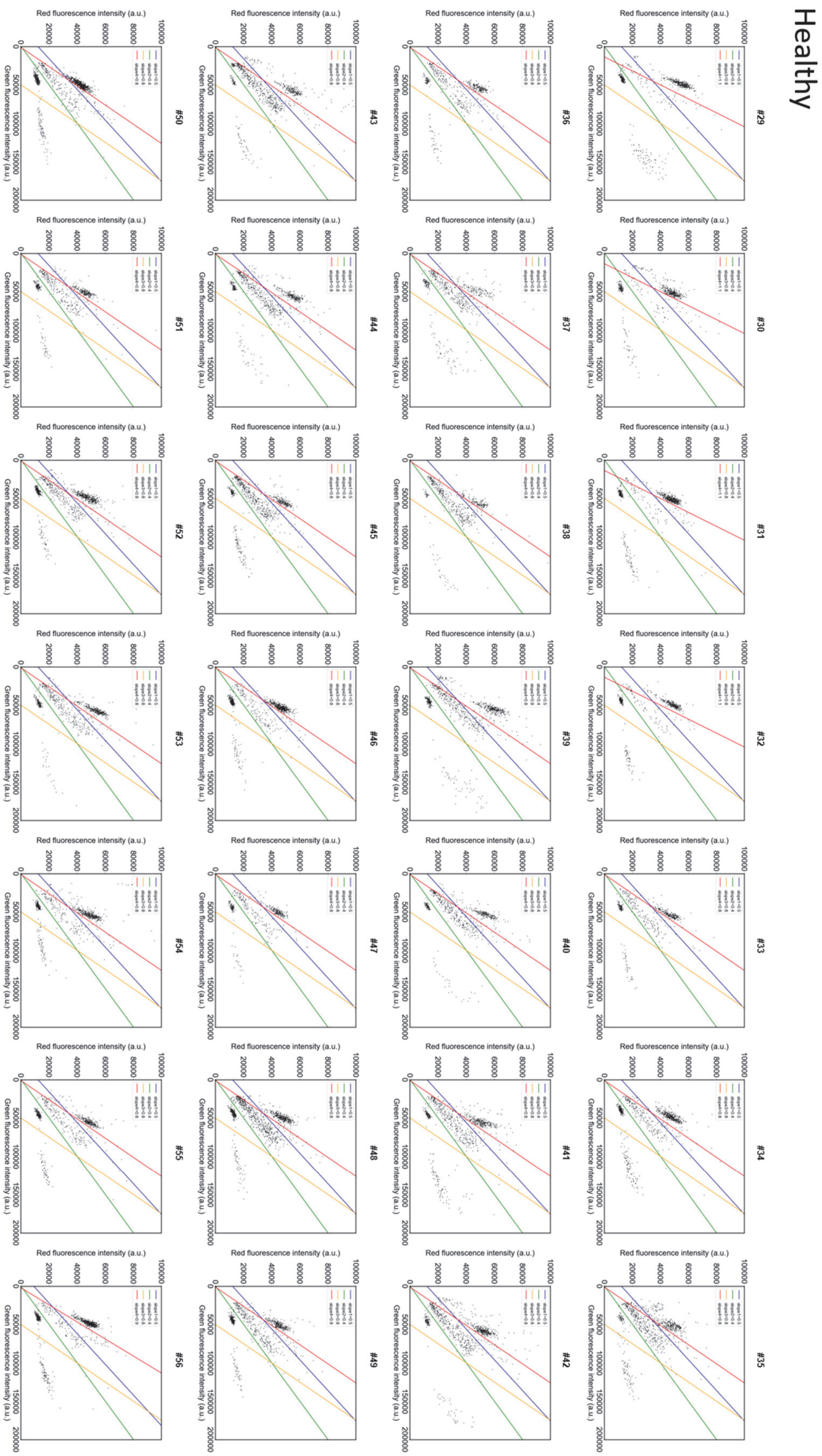


**Figure S7.** Scattered plot of single-molecule CPA activities of healthy subjects (#29-#56)

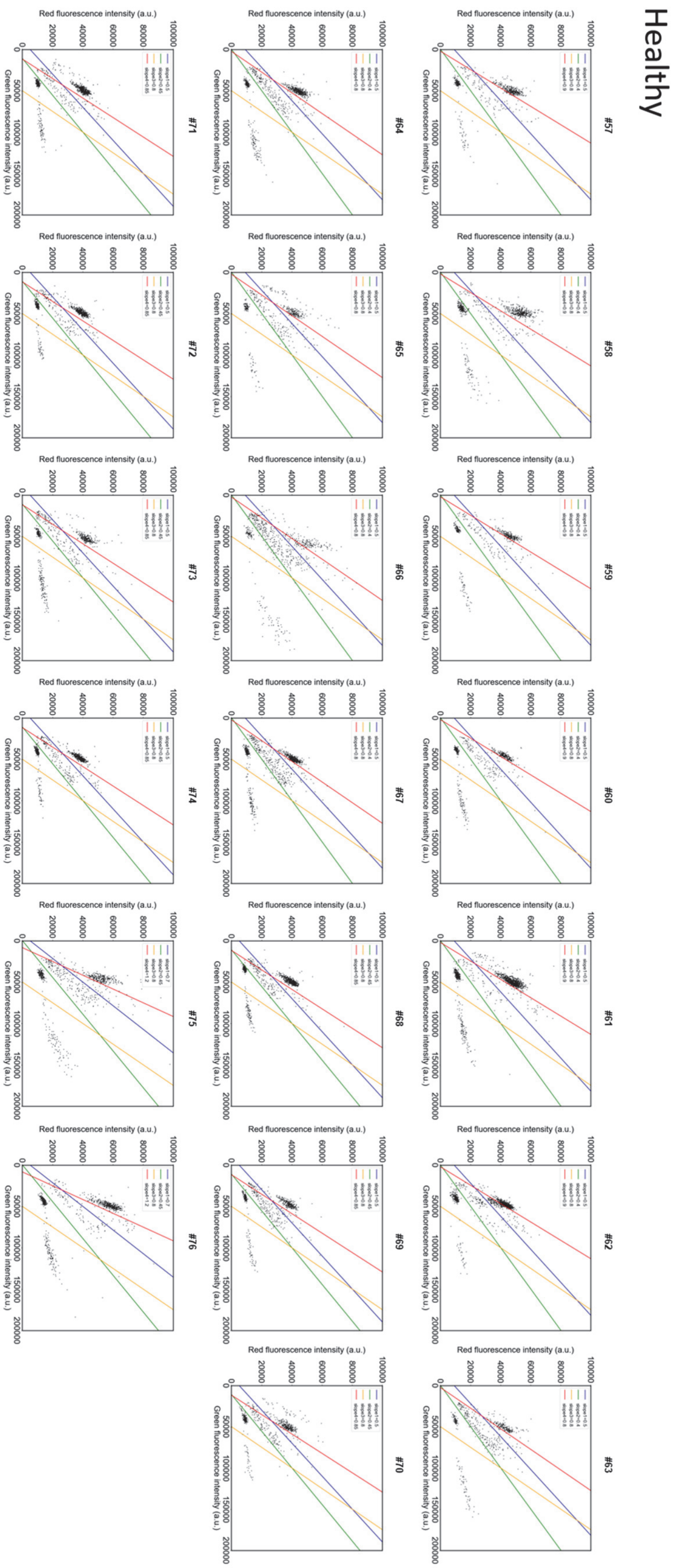


**Figure S8.** Scattered plot of single-molecule CPA activities of healthy subjects (#57-#76)

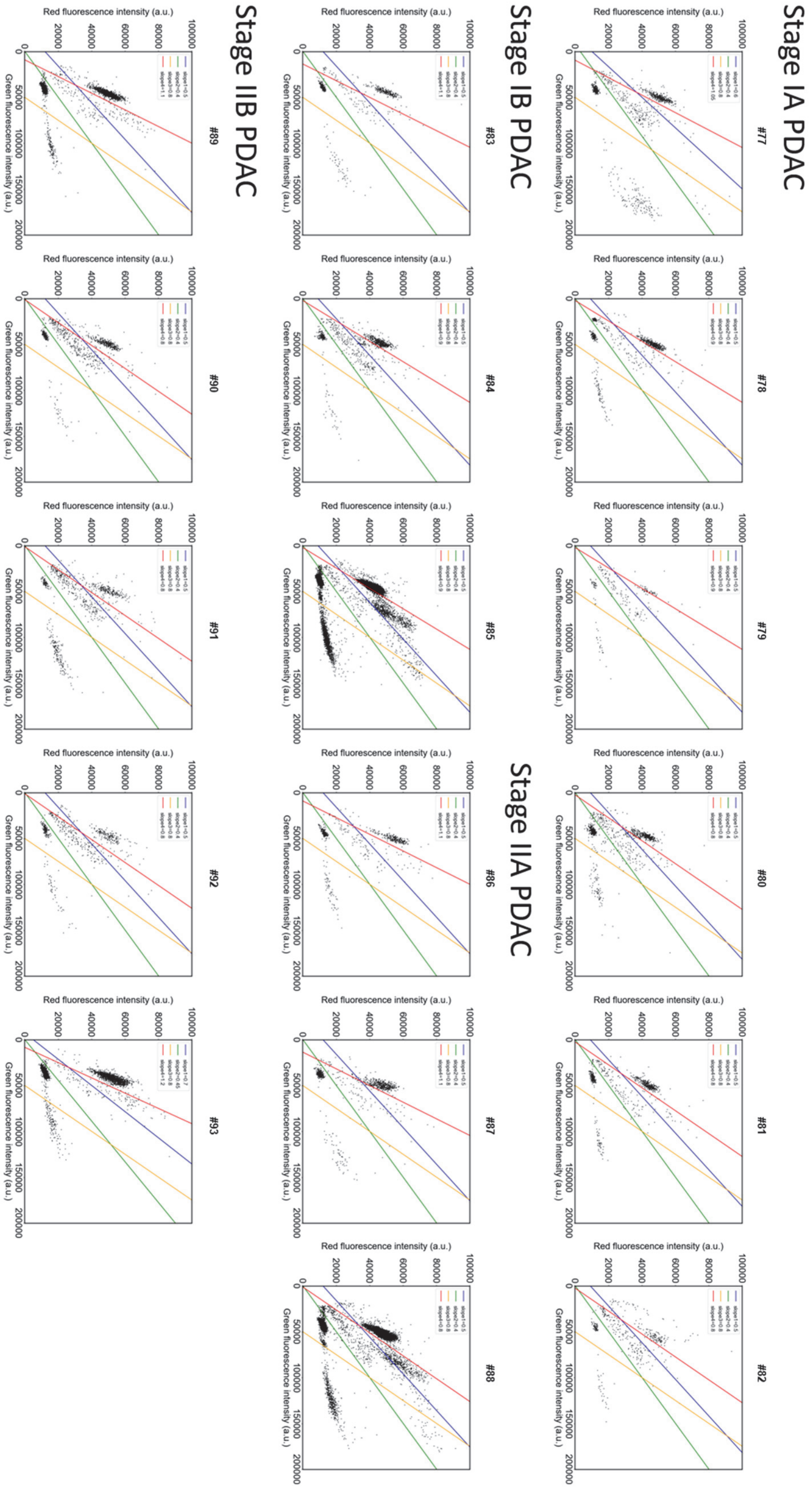


**Figure S9.** Scattered plot of single-molecule CPA activities of pancreatic cancer patients (stage I-II PDAC).

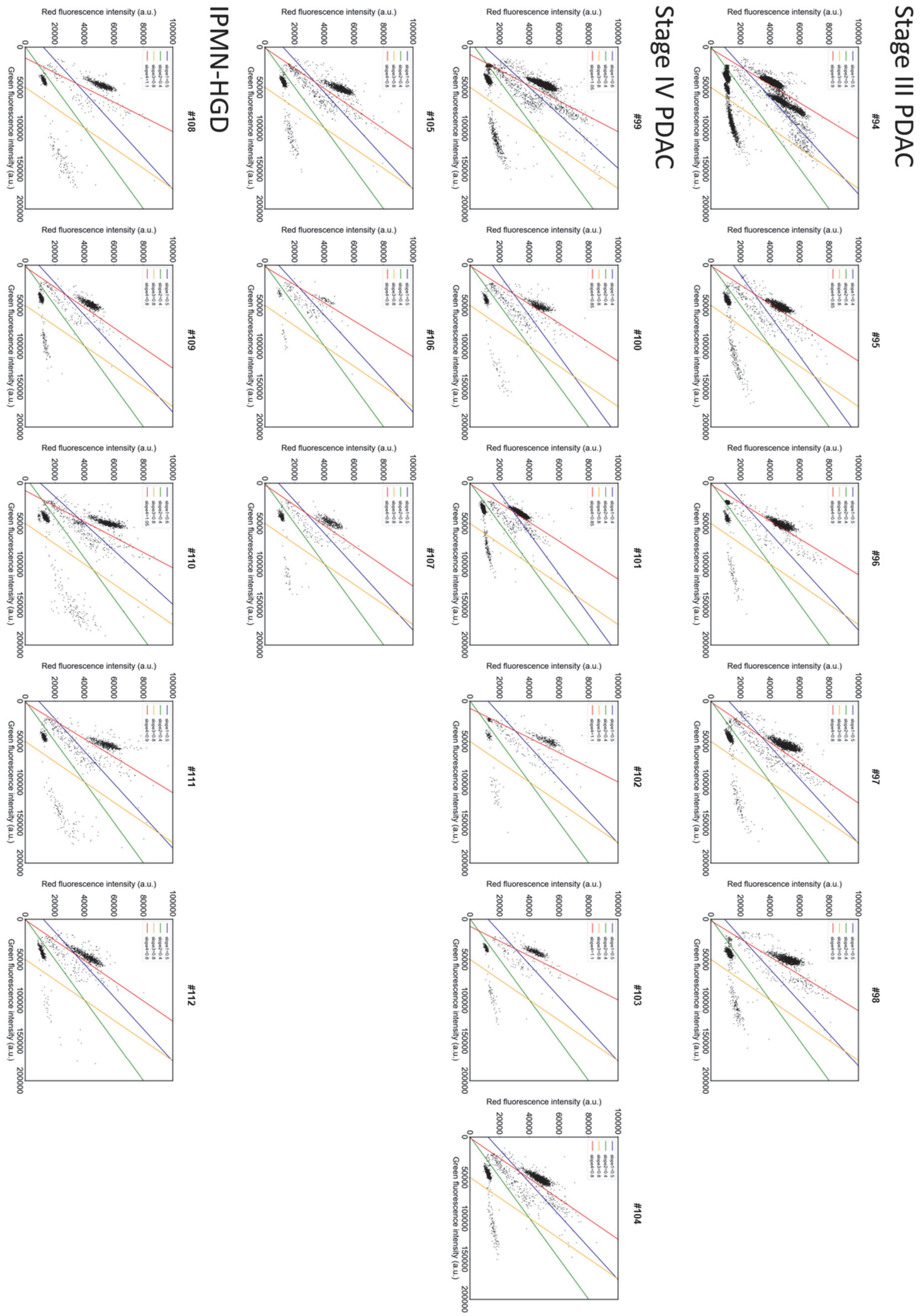


**Figure S10.** Scattered plot of single-molecule CPA activities of pancreatic cancer patients (stage III-IV PDAC and malignant IPMN).

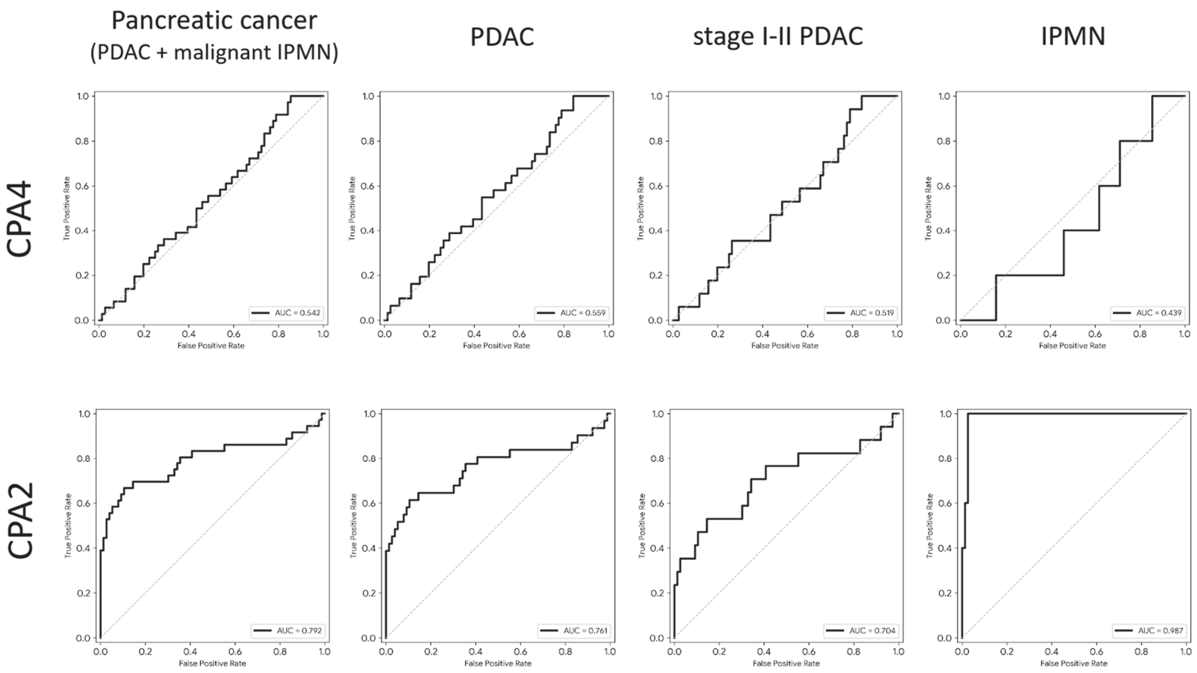


**Figure S11.** ROC curves of CPA4 and CPA2 for various types of pancreatic cancer.

**Supplementary data for synthesis and characterization of compounds**

**Stereochemical considerations of ProTide probes**

Due to the presence of both the chiral phosphorus center in the phosphate/phosphonate moiety and the stereogenic centers in the peptide, the resulting ProTide probes are, in principle, obtained as mixtures of diastereomers. In the present study, these diastereomers were not separated and were treated collectively in all experiments.

**Nomenclature of ProTide-based fluorogenic probes**

ProTide-based fluorogenic probes described in this study are named using a modular notation that reflects their molecular architecture. The general format is:

*[Fluorophore]–[Linker]–[ProTide type]–[Peptide sequence]*

where each element corresponds to a distinct structural component of the probe. The fluorophore is indicated by an abbreviated name, such as sTG for sulfonated TokyoGreen[2] and Res for resorufin. When present, a self-immolative *p*-hydroxybenzyl alcohol linker is denoted as PHBA. The ProTide moiety is classified based on the nature of the phosphorus center. Ethyl phosphoramidate–based ProTides are designated as P(Et), whereas *tert*-butyl phosphonamidate–based ProTides are designated as PA(*t*Bu), reflecting their phosphonate character. The peptide sequence corresponds to the C-terminal dipeptide substrate and is written using the standard one-letter amino acid code, with the P1 residue listed first followed by the P1' residue.
In a previous report[3], ProTide-based fluorogenic probes were referred to as eaSoul and tSoul. In the present study, eaSoul corresponds to the phosphate-type ProTide denoted as P(Et), while tSoul corresponds to the phosphonate-type ProTide denoted as PA(*t*Bu). The updated nomenclature is used throughout this manuscript for clarity and consistency.

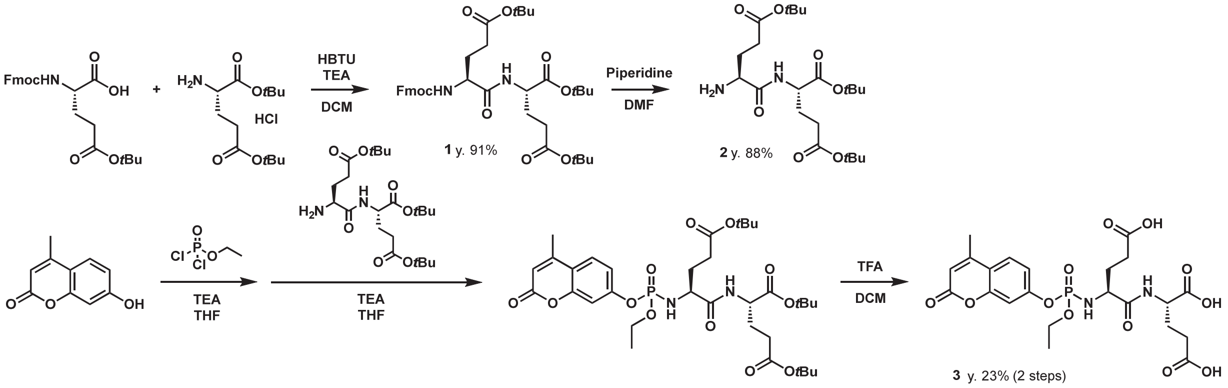


**Scheme S1.** Preparation of 4MU-P(Et)-EE using liquid phase synthesis.

**Preparation of Fmoc-Glu(O*t*Bu)-Glu-(O*t*Bu)-O*t*Bu (1)**

Fmoc-Glu(O*t*Bu)-OH (790 mg, 1.9 mmol), $NH_2$-Glu(O*t*Bu)-O*t*Bu HCl salt (515 mg, 1.7 mmol) and HBTU (770 mg, 2.0 mmol) were dissolved in dichloromethane (DCM, 25 mL), and triethylamine (TEA, 700 μL, 5.1 mmol) was added. After stirring the reaction at 25°C for 4 h, The solution was washed with sat. $NaHCO_3$ *aq*. The organic layer was dried over $Na_2SO_4$, filtered, and evaporated. The product was purified using column chromatography (silica; AcOEt-Hexane). **1** (1062 mg, y. 91%) was acquired as colorless solid.

$^1$H-NMR (400 MHz, DMSO-$d_6$) δ 8.20 (d, 1H, *J* = 8.4 Hz), 7.85 (d, 2H, *J* = 8.4 Hz), 7.72 (t, 2H, *J* = 8.4 Hz), 7.52 (d, 1H, *J* = 8.4 Hz), 7.38 (t, 2H, *J* = 8.4 Hz), 7.29 (t, 2H, *J* = 8.4 Hz), 3.9-4.2 (m, 5H), 2.29 (m, 4H), 1.8-2.0 (m, 4H), 1.35 (s, 9H), 1.34 (s, 9H), 1.34 (s, 9H).

$^{13}$C-NMR (100 MHz, $CDCl_3$) δ 172.2, 172.0, 171.3, 156.4, 144.4, 144.3, 141.2, 128.2, 127.6, 125.9, 120.7, 81.2, 80.3, 80.2, 66.2, 52.8, 52.3, 47.2, 31.9, 31.4, 28.3, 28.2, 28.1, 27.9, 26.6.

LRMS (ESI$^+$): $m/z$ = 667.3 $(M+H)^+$

**Preparation of $NH_2$-Glu(O*t*Bu)-Glu-(O*t*Bu)-O*t*Bu (2)**

**1** (1043 mg, 1.56 mmol) was dissolved in piperidine (5 mL) and *N*,*N*-dimethylformamide (DMF, 20 mL), and the reaction was stirred at 25°C for 60 min. $H_2O$ was added, and the product was extracted with AcOEt three times. The combined organic layer was washed with brine, dried over $Na_2SO_4$, filtered, and evaporated. The product was purified using column chromatography (amino; AcOEt-Hexane). **2** (316 mg, y. 88%) was acquired as colorless solid.

$^1$H-NMR (400 MHz, DMSO-$d_6$) δ 8.04 (d, 1H, *J* = 8.4 Hz), 4.09 (m, 1H), 3.12 (m, 1H), 2.20 (m, 4H), 1.6-1.8 (m, 4H), 1.35 (s, 9H), 1.34 (s, 9H), 1.34 (s, 9H).

LRMS (ESI$^+$): $m/z$ = 445.3 $(M+H)^+$

**Preparation of 4MU-P(Et)-EE (3)**

Ethyl phosphorodichloridate (50 μL, 0.42 mmol) was dissolved in dry THF (2 mL) under an Ar atmosphere at -78°C. 4-Methylumberiferrone (4MU, 49 mg, 0.28 mmol) dissolved in dry THF (2 mL) and TEA (90 μL, 0.65 mmol) were added successively into the solution dropwise, and the reaction was stirred at -78°C for 15 min. 2 (186 mg, 0.42 mmol) and TEA (160 μL, 1.15 mmol) in dry THF (2 mL) was added dropwise. The reaction was stirred at -78°C for 35 min. To the reaction, sat. NH4Cl aq. was added, and the product was extracted using AcOEt three times. The combined organic layer was washed with brine, dried over $Na_2SO_4$, filtered, and evaporated. The protected product was purified using column chromatography (amino; AcOEt-Hexane). The deprotection was performed by dissolving the purified product in dry acetonitrile (1 mL) and trifluoroacetic acid (TFA, 1 mL) and stirring at 25°C for 40 min. The product was purified over prep. HPLC ($C_{18}$, A: 0.1% TFA in $H_2O$, B = 80%-AcCN-20% $H_2O$-0.1% TFA, A:B = 80:20 to 0:100, 50 min). **3** (35.1 mg, y. 23%, 2 steps) was acquired as colorless solid.

$^1$H-NMR (400 MHz, $CD_3OD$) δ 8.30 (m, 1H), 7.76 (m, 1H), 7.24 (m, 2H), 6.26 (s, 1H), 4.37 (m, 1H), 4.21 (m, 2H), 3.86 (m, 1H), 2.45 (s, 3H), 2.3-2.4 (m, 4H), 1.8-2.2 (m, 4H), 1.36 (m, 3H).

LRMS (ESI$^+$): $m/z$ = 543.0 $(M+H)^+$

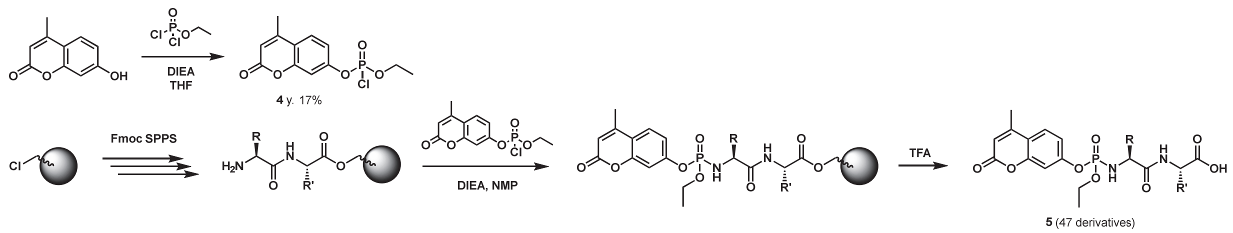

**Scheme S2.** Solid-phase synthesis of ProTide probes.

**Preparation of phosphorochloridate of 4MU (4)**

Ethyl phosphorodichloridate (400 μL, 3 mmol) was dissolved in dry THF (3 mL) under an Ar atmosphere at 0°C. 4-Methylumberiferrone (400 mg, 2.3 mmol) dissolved in *N*-methylpyrrolidone (NMP, 2 mL) and *N*,*N*-diisopropylethylamine (DIEA, 1 mL) was added into the solution dropwise, and the reaction was stirred at 25°C for 10 min. The reaction was loaded onto silica gel column (silica) for purification using prep. MPLC (AcOEt : Hexane = 80:20 to 0:100, 10 min). The fractions containing the desired product were collected and evaporated. The product (120 mg, y. 17%) was stored at -20°C.

$^1$H-NMR (400 MHz, $CDCl_3$) δ 7.63 (d, 1H, *J* = 8.4 Hz), 7.2-7.3 (m, 2H), 6.30 (s, 1H), 4.45 (m, 2H), 2.45 (s, 3H), 1.50 (t, 3H, *J* = 7.2 Hz).

$^{13}$C-NMR (100 MHz, $CDCl_3$) δ 160.3, 154.4, 151.8, 126.2, 118.3, 116.8, 116.7, 114.9, 109.4, 67.5, 18.9, 15.9.

LRMS (ESI$^+$): *m/z* = 303.1 $(M+H)^+$

**Preparation of ProTide-based 4MU probe by solid phase synthesis (5).**

For loading of first amino acid on resin, 2-chlorotrityl chloride (CTC) resin (60 mg) was mixed with $N_\alpha$-Fmoc-protected amino acid (0.4 M in DMF, 800 μL), DIEA (400 μL) and DMF (800 μL) and stirred at 25°C for 4 h. Then, the solution was removed, and the resin was mixed with Fmoc-AA (0.4 M in DMF, 800 μL), DIEA (400 μL) and DMF (800 μL) and stirred at 25°C for and stirred at 25°C for 2 h. The resin was washed six times with DMF. For elongation of peptide, Fmoc deprotection and coupling were performed sequentially until the peptide with desired lengths were acquired. For Fmoc deprotection, piperidine (40% in DMF; 1.2 mL) was added to beads and stirred at 25°C for 3 min. After removing the solution, piperidine (40% in DMF; 600 μL) and DMF (600 μL) were added and stirred for 12 min. Then, the resin was washed six times with DMF. For peptide elongation, Fmoc-AA (0.4 M in DMF, 800 μL), HATU (0.4 M in DMF, 840 μL), DIEA (1.6 M in NMP, 400 μL) were added to resin and the reaction was stirred at 30°C for 40 min. Resin was washed three times with DMF. To the resin loaded with the desired peptide, NMP (200 μL) and DIEA (200 μL) were added, and **5** (15 mg, 50 μmol) dissolved in NMP (200 μL) was added. The reaction was stirred at 25°C for 2 h, and the resin was washed three times with DMF and three times with DCM. The ProTide probe was cleaved from resin by treating with 90% trifluoroacetic acid (TFA)-10% $H_2O$ (5 mL) for 10 min. The reaction was diluted with $H_2O$ (5 mL) and the purity of the product was checked using LC-MS. If the sufficient purity (> 90% at 320 nm) was confirmed, the solution was fresh frozen using liquid $N_2$ and freeze-dried. If the purification was required, the solution was directly injected into ODS column for purification over prep. MPLC ($C_{18}$, A: 0.1% TFA in $H_2O$,

B = 0.1% TFA in AcCN, A:B = 99:1 to 0:100, 15 min). The fractions containing the desired product were collected, evaporated to remove AcCN, fresh frozen using liquid $N_2$, and freeze-dried. The DMSO stock was prepared, and the concentration was adjusted to 10 mM by measuring the absorbance of the diluted solution in sodium phosphate buffer ($\varepsilon$ = 12,000, 320 nm[4]).

**4MU-based ProTide probes**

LC Chromatogram was monitored at 320 nm ($H_2O$-0.1% TFA/AcCN-0.1% TFA = 95/5 to 0/100, 3.5 min). The chromatograms were background subtracted using the blank data (injecting DMSO). The ESI-MS ($ESI^+$) spectra of the main peak are shown.

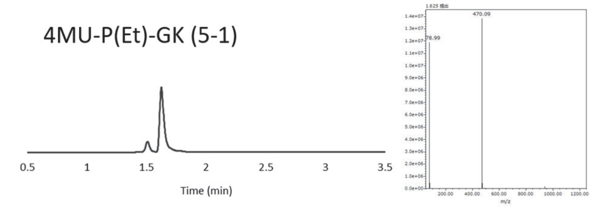


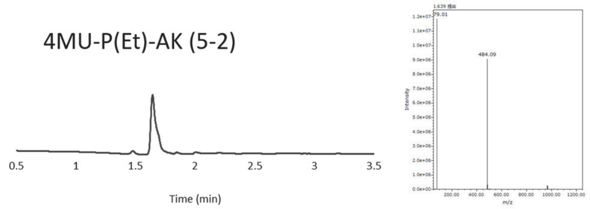


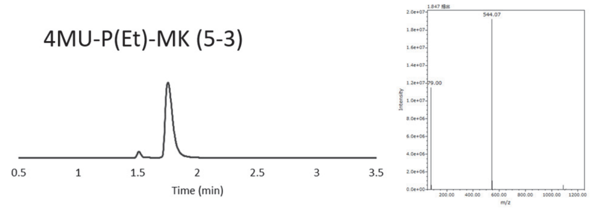


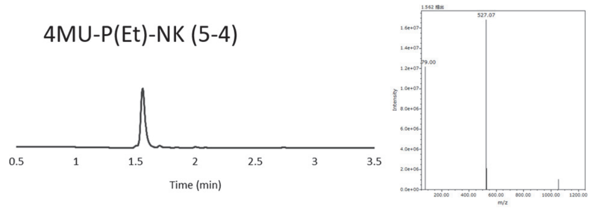


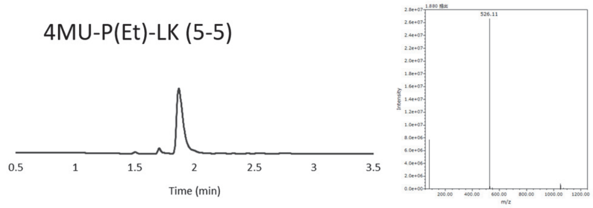


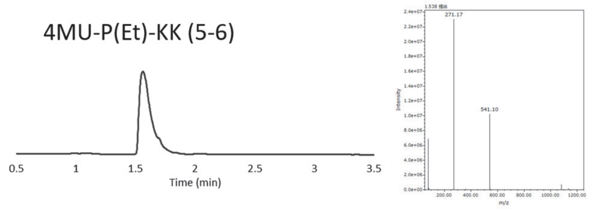


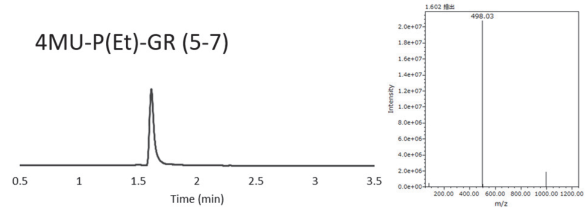


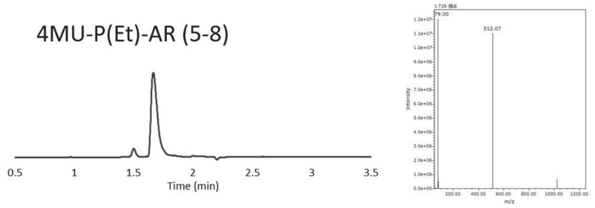


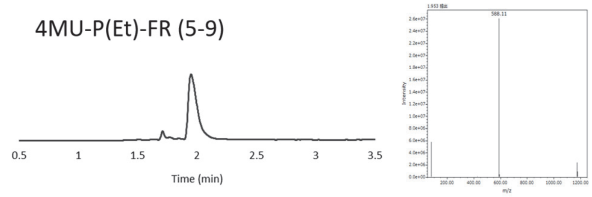


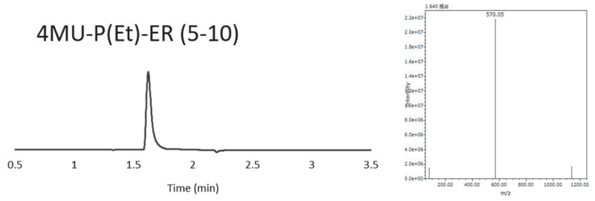


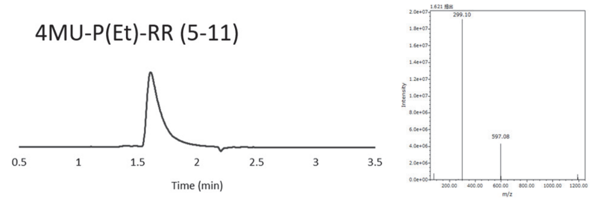


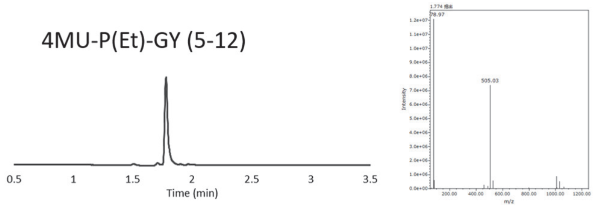

4MU-P(Et)-AY (5-13)

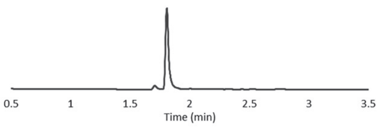


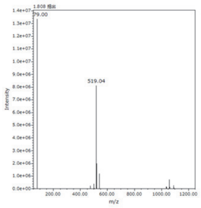


4MU-P(Et)-GF (5-14)

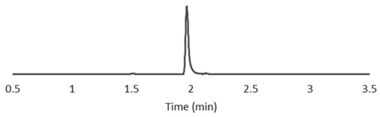


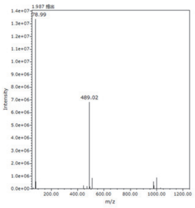


4MU-P(Et)-PF (5-15)

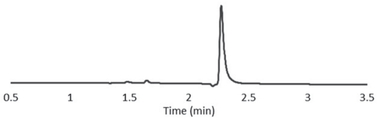


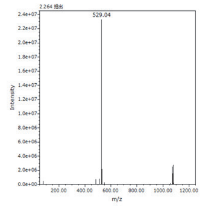


4MU-P(Et)-AH (5-16)

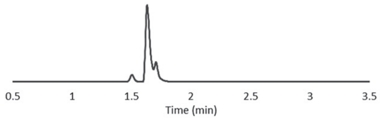


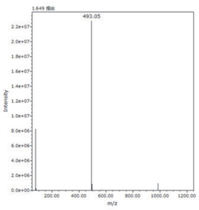


4MU-P(Et)-FH (5-17)

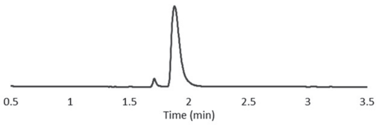


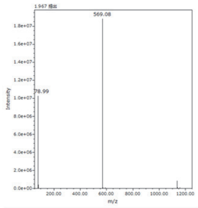


4MU-P(Et)-DL (5-18)

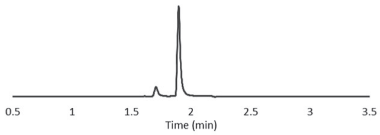


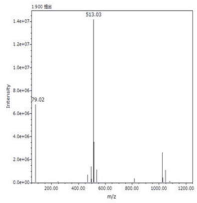


4MU-P(Et)-AL (5-19)

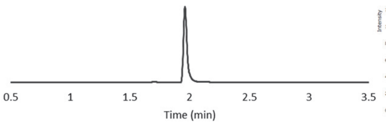


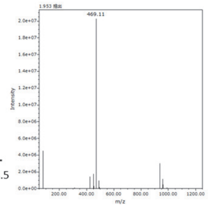


4MU-P(Et)-VL (5-20)

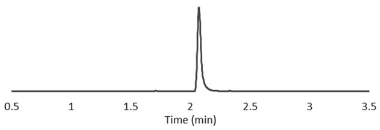


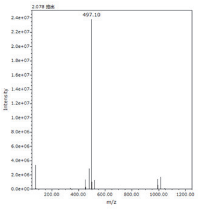


4MU-P(Et)-FL (5-21)

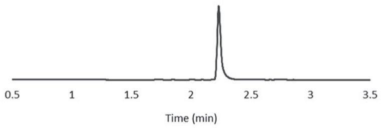


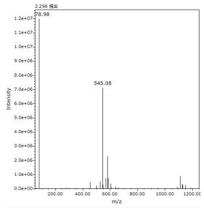


4MU-P(Et)-HL (5-22)

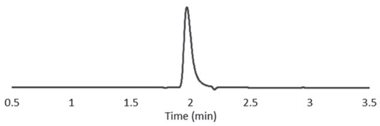


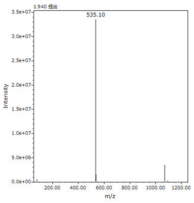


4MU-P(Et)-AI (5-23)

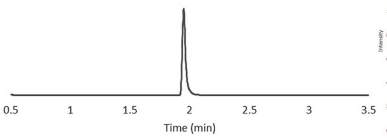


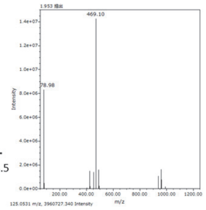


4MU-P(Et)-AV (5-24)

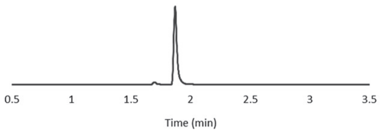


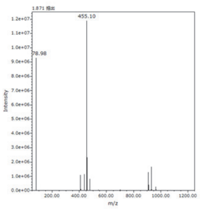

4MU-P(Et)-AS (5-25)

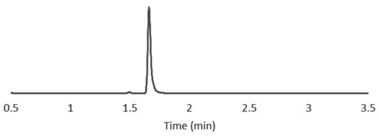


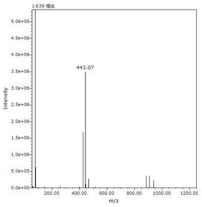


4MU-P(Et)-DS (5-26)

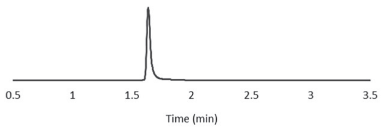


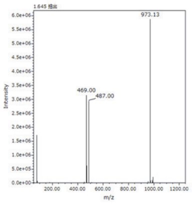


4MU-P(Et)-AT (5-27)

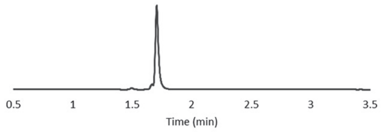


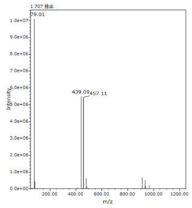


4MU-P(Et)-VT (5-28)

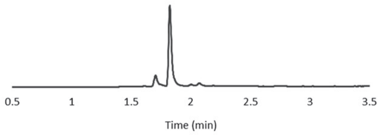


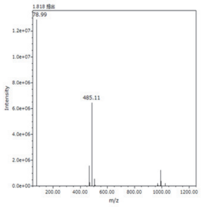


4MU-P(Et)-NT (5-29)

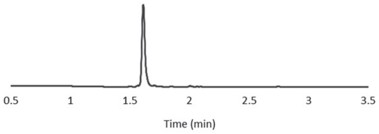


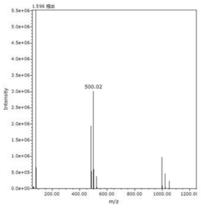


4MU-P(Et)-GQ (5-30)

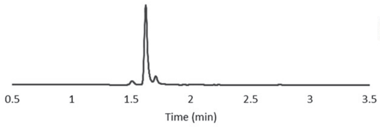


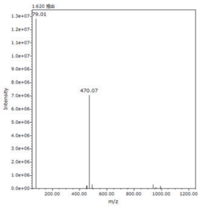


4MU-P(Et)-AQ (5-31)

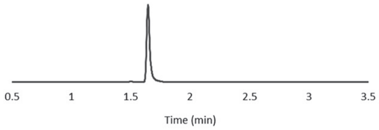


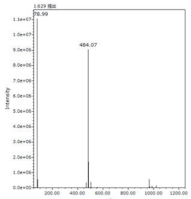


4MU-P(Et)-GE (5-32)

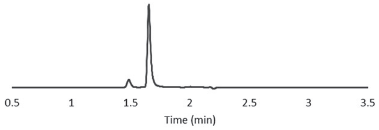


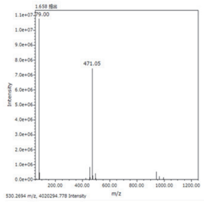


4MU-P(Et)-AE (5-33)

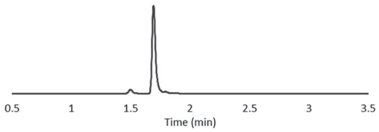


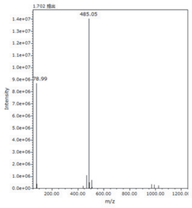


4MU-P(Et)-FE (5-34)

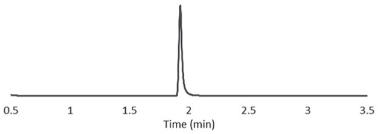


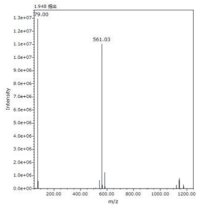


4MU-P(Et)-EE (5-35)

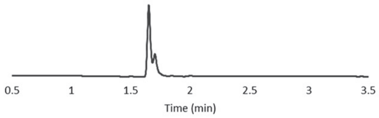


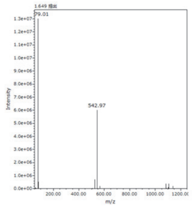


4MU-P(Et)-RE (5-36)

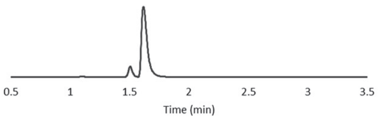


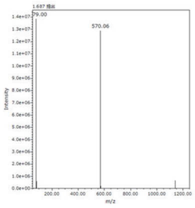

4MU-P(Et)-VA (5-37)

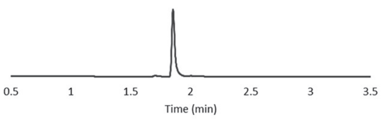


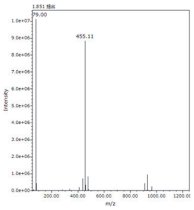


4MU-P(Et)-AM (5-38)

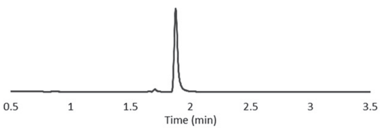


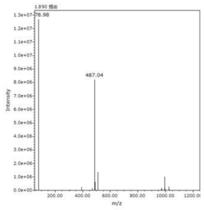


4MU-P(Et)-FM (5-39)

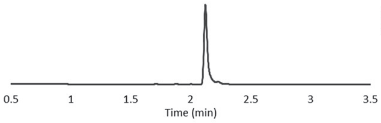


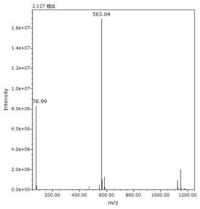


4MU-P(Et)-AP (5-40)

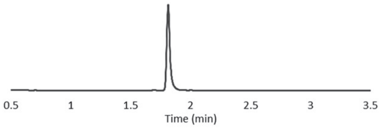


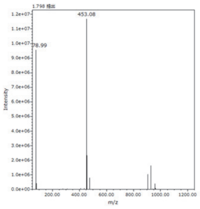


4MU-P(Et)-HP (5-41)

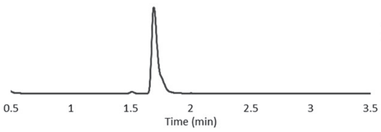


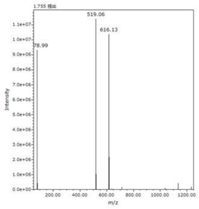


4MU-P(Et)-AG (5-42)

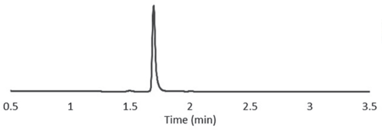


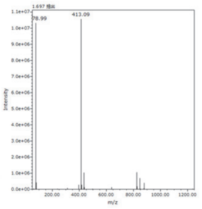


4MU-P(Et)-LG (5-43)

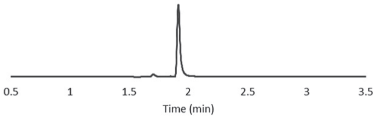


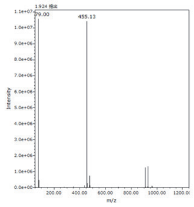


4MU-P(Et)-RG (5-44)

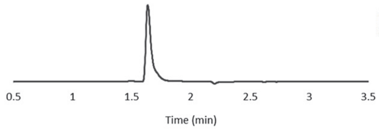


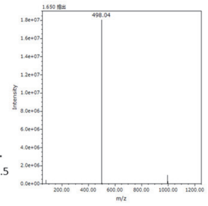


4MU-P(Et)-T(O*t*Bu)A (5-45)

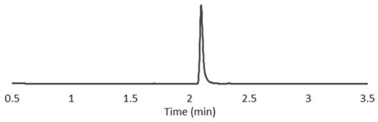


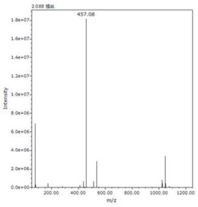


4MU-P(Et)-GR(Pbf) (5-46)

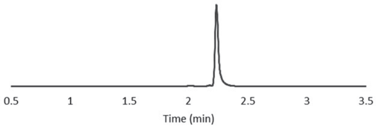


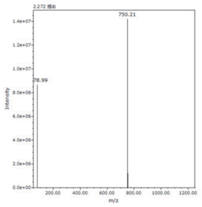


4MU-P(Et)-ER(Pbf) (5-47)

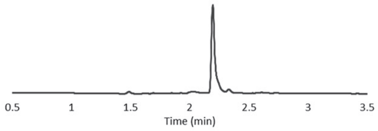


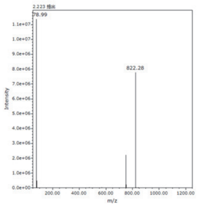

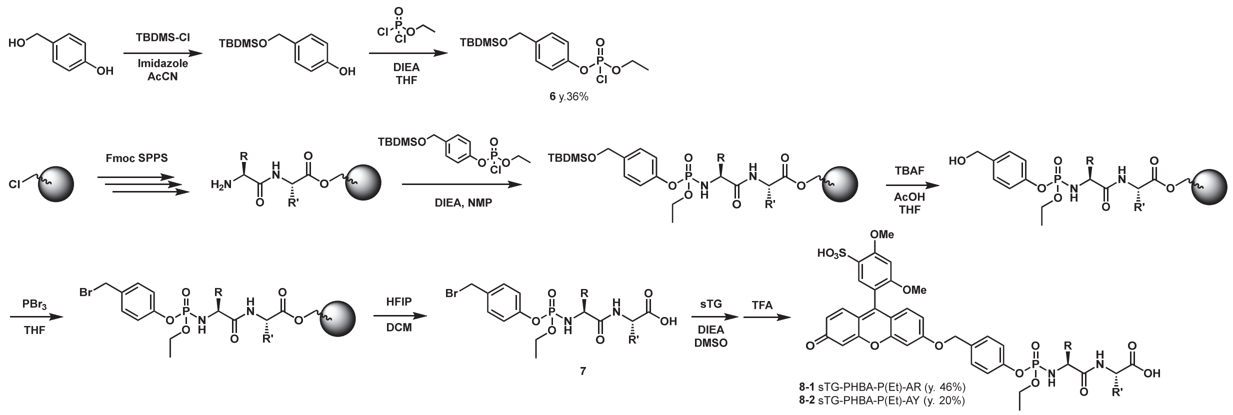


**Scheme S3.** Preparation of single-molecule enzyme activity assay probe.

**Preparation of phosphorodichloridate of PHBA-TBDMS (6)**

*p*-Hydroxybenzyl alcohol (PHBA) TBDMS was prepared according to the literature[3]. Ethyl phosphorodichloridate (600 μL, 5.0 mmol) was dissolved in dry THF (3 mL) under an Ar atmosphere at 0°C. PHBA TBDMS (200 mg, 0.42 mmol) dissolved in THF (2 mL) and DIEA (500 μL) was added into the solution dropwise, and the reaction was stirred at 25°C for 15 h. The reaction was loaded onto silica gel column (silica) for purification using prep. MPLC (AcOEt : Hexane = 80:20 to 0:100, 10 min). The fractions containing the desired product were collected and evaporated. The product (55 mg, y. 36%) was used for the next reaction immediately.

$^{1}$H-NMR (400 MHz, $CDCl_3$) δ 7.33 (d, 2H, *J* = 8.4 Hz), 7.23 (d, 2H, *J* = 8.4 Hz), 4.72 (s, 2H), 4.40 (m, 2H), 1.47 (t, 3H, *J* = 7.2 Hz), 0.94 (s, 9H), 0.10 (s, 6H).

$^{13}$C-NMR (100 MHz, $CDCl_3$) δ 148.7, 139.6, 127.6, 120.2, 66.9, 64.3, 26.0, 25.8, 18.5, -5.1.

LRMS (ESI$^+$): *m/z* = 361.2 $(M+H)^+$ for sample treated in methanol for 5 min.

**Preparation of Bromide (7) using solid phase syntehsis**

Preparation of peptide on CTC resin was performed in the same procedure as in **2**. Bromide was prepared following the below procedures. At each step, the progress of the reaction was confirmed by cleaving off the small amount of sample using 20% 1,1,1,3,3,3-hexafluoropropan-2-ol (HFIP)-80% DCM.

**ProTide formation**

To the resin loaded with the desired peptide, NMP (200 μL) and DIEA (200 μL) were added, and **7** (36 mg, 100 μmol) dissolved in NMP (200 μL) was added. The reaction was stirred at 25°C for 2 h, and the resin was washed three times with DMF and three times with DCM.

**TBDMS deprotection**

THF (1.5 mL) mixed with TBAF (1 M in THF, 500 μL) and acetic acid (40 μL) was added to resin, and the mixture was stirred at 25°C for 3 h. The resin was washed three times with DMF and three times with DCM. After the reaction, if the free peptide was detected, they were capped by treating with *N*-succinimidyl acetate (157 mg, 1.0 mmol) dissolved in DMF (2 mL) and DIEA (500 μL) for 1 h.

**Bromination**

Dry THF (2 mL) was added to resin, and the mixture was stirred at 25°C for 20 min. While stirring, $PBr_3$ (50 µL, 0.53 mmol) was added dropwise, and the mixture was stirred for 1 min. The solution was removed, and the resin was washed three times with DMF, once with MeOH, then three times with DCM.

**Release and purification**

The resin was treated with 20% HFIP-80% DCM (1 mL) at 25°C for 30 min, and the solution was collected. Resin was washed with DCM (500 µL) and AcCN (3 mL), and all solutions were combined and concentrated *in vacuo*. The remaining was dissolved in $H_2O$ and was purified over prep. MPLC ($C_{18}$, A: 0.1% TFA in $H_2O$, B = 0.1% TFA in AcCN, A:B = 99:1 to 0:100, 15 min). The fractions containing the desired product were combined, and immediately concentrated *in vacuo*. Within 30 min, the solution was fresh frozen using liq. $N_2$ and freeze-dried.

**Preparation of sTG-based ProTide-based probes (8)**

sTG[2] (4.3 mg, 10 µmol) was dissolved in DMSO (15 µL) and DIEA (10 µL) and DCM (500 µL) was added. The solution was added to prepared bromide (5 µmol) in 1.5 mL plastic tube, and DCM was removed using centrifugal evaporator (EYELA; CVE-2200) at 25-30°C. The concentrated solution was stirred at 25°C for 12 h. After confirming the product formation using LC-MS, 90% TFA-10% $H_2O$ (500 µL) was added, and the mixture was stirred at 25°C for 30 min. After confirming the product formation using LC-MS, the reaction was diluted with $H_2O$ and was purified over prep. MPLC ($C_{18}$, A: 0.1% TFA in $H_2O$, B = 0.1% TFA in AcCN, A:B = 99:1 to 0:100, 15 min). The fractions containing the desired product were combined, and immediately concentrated *in vacuo*. Then, the compound was purified again over prep. MPLC ($C_{18}$, A: 10 mM triethylamine acetate in $H_2O$, B = AcCN, A:B = 99:1 to 0:100, 15 min). The fractions containing the desired product were combined, $H_2O$ was added, and the solution was fresh frozen using liq. $N_2$ and freeze-dried. The DMSO stock was prepared, and the concentration was adjusted to 10 mM by measuring the absorbance of the diluted solution in sodium phosphate buffer (ε = 30,000, 450 nm[2], y. 46% for **8-1**, 20% for **8-2**).

**sTG-PHBA-P(Et)-AR (8-1)**

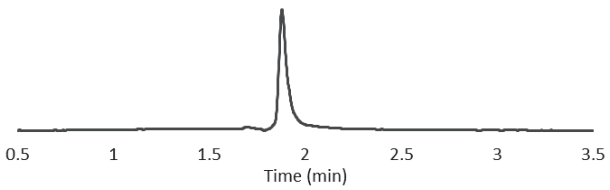


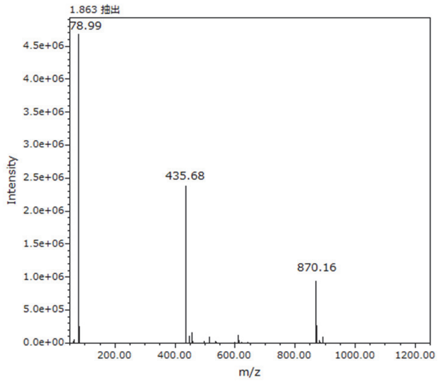


LC Chromatogram was monitored at 450 nm ($H_2O$-0.1% TFA/AcCN-0.1% TFA = 95/5 to 0/100, 3.5 min).

LRMS (ESI$^+$): *m/z* = 435.7 $(M+2H)^{2+}$, 870.2 $(M+H)^+$

**sTG-PHBA-P(Et)-AY (8-2)**

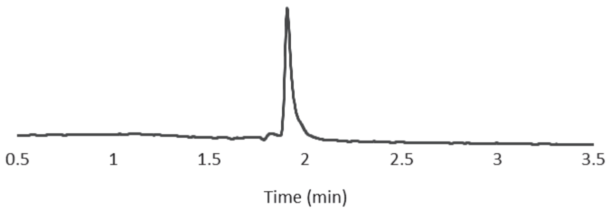


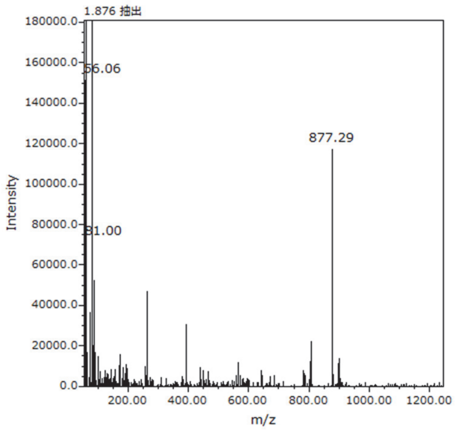


LC Chromatogram was monitored at 450 nm ($H_2O$-0.1% TFA/AcCN-0.1% TFA = 95/5 to 0/100, 3.5 min).
LRMS (ESI$^+$): $m/z$ = 877.3 $(M+H)^+$

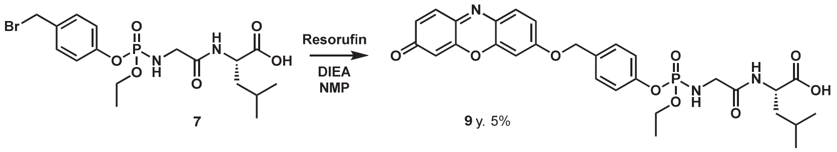


**Scheme S4.** Preparation of Res-PHBA-P(Et)-GL.

### Preparation of Res-PHBA-P(Et)-GL (9)

Resorufin (6.8 mg, 32 μmol) was dissolved in NMP (15 μL) and DIEA (10 μL) and DCM (500 μL) was added. The solution was added to prepared bromide (10 μmol) in 1.5 mL plastic tube, and DCM was removed using centrifugal evaporator (EYELA; CVE-2200) at 25-30°C. The concentrated solution was stirred at 25°C for 3 h. After confirming the product formation using LC-MS, 90% TFA-10% $H_2O$ (500 μL) was added, and the mixture was stirred at 25°C for 30 min. After confirming the product formation using LC-MS, the reaction was diluted with $H_2O$ and was purified over prep. MPLC ($C_{18}$, A: 0.1% TFA in $H_2O$, B = 0.1% TFA in AcCN, A:B = 99:1 to 0:100, 15 min). The fractions containing the desired product were combined, and immediately concentrated *in vacuo*. Then, the compound was purified again over prep. MPLC ($C_{18}$, A: 10 mM triethylamine acetate in $H_2O$, B = AcCN, A:B = 99:1 to 0:100, 15 min). The fractions containing the desired product were combined, $H_2O$ was added, and the solution was fresh frozen using liq. $N_2$ and freeze-dried. The DMSO stock was prepared, and the concentration was adjusted to 10 mM by measuring the absorbance of the diluted solution in sodium phosphate buffer (ε = 13,200, 450 nm[5], y. 5%).

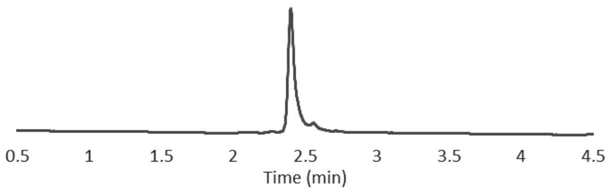


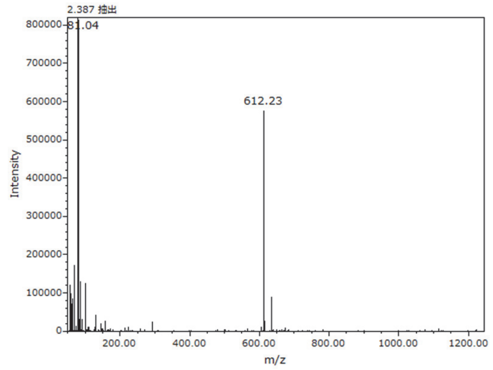


LC Chromatogram was monitored at 450 nm ($H_2O$-0.1% TFA/AcCN-0.1% TFA = 95/5 to 0/100, 3.5 min).

LRMS (ESI$^+$): *m/z* = 612.2 (M+H)$^+$

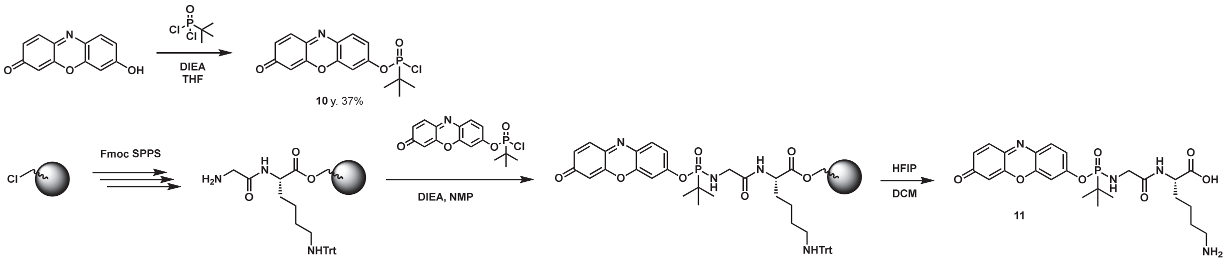


**Scheme S5.** Preparation of Res-PA(*t*Bu)-GK.

**Preparation of phosphonodichloridate of resorufin (10)**

*tert*-Butylphosphonic dichloride (220 mg, 1.3 mmol) was dissolved in dry THF (4 mL) under an Ar atmosphere at 0°C. Resorufin (90 mg, 0.42 mmol) dissolved in NMP (2 mL) and DIEA (400 μL) was added into the solution dropwise, and the reaction was stirred at 25°C for 18 h. The reaction was loaded onto silica gel column (silica) for purification using prep. MPLC (AcOEt : Hexane = 80:20 to 0:100, 10 min). The fractions containing the desired product were collected and evaporated. The product (55 mg, y. 37%) was stored at -20°C.

$^1$H-NMR (400 MHz, $CDCl_3$) δ 7.78 (d, 1H, *J* = 8.4 Hz), 7.42 (d, 1H, *J* = 8.4 Hz), 7.2-7.3 (m, 2H), 6.86 (d, 1H, *J* = 8.4 Hz), 6.32 (s, 1H), 1.46 (d, 9H, *J* = 15.1 Hz).

$^{13}$C-NMR (100 MHz, $CDCl_3$) δ 186.3, 152.4, 149.3, 148.5, 144.7, 135.4, 134.9, 131.8, 131.2, 118.4, 108.9, 107.5, 39.4, 38.2, 24.4

LRMS (ESI$^+$): *m/z* = 352.0 (M+H)$^+$

**Preparation of Res-PA(*t*Bu)-GK (11)**

$NH_2$-Gly-Lys(Trt)-OH was prepared on CTC resin following the same procedure as in **2**. **11** (18 mg, 50 μmol) dissolved in NMP (500 μL) and DIEA (400 μL) was added to resin and the mixture was stirred at 25°C for 12 h. Resin was washed three times with DMF and three times with DCM. The resin was treated with 20% HFIP-80% DCM (1 mL) at 25°C for 30 min, and the solution was collected. Resin was washed with DCM (500 μL) and AcCN (3 mL), and all solutions were combined and concentrated *in vacuo*. The remaining was dissolved in $H_2O$ and was purified over prep. MPLC ($C_{18}$, A: 0.1% TFA in $H_2O$, B = 0.1% TFA in AcCN, A:B = 99:1 to 0:100, 15 min). The fractions containing the desired product were combined, and immediately concentrated *in vacuo*. Then, the compound was purified again over prep. MPLC ($C_{18}$, A: 10 mM triethylamine acetate in $H_2O$, B = AcCN, A:B = 99:1 to 0:100, 15 min). The fractions containing the desired product were combined, $H_2O$ was added, and the solution was fresh frozen using liq. $N_2$ and freeze-dried. The DMSO stock was prepared, and the concentration was adjusted to 10 mM by measuring the absorbance of the diluted solution in sodium phosphate buffer (ε = 13,200, 450 nm[5]).

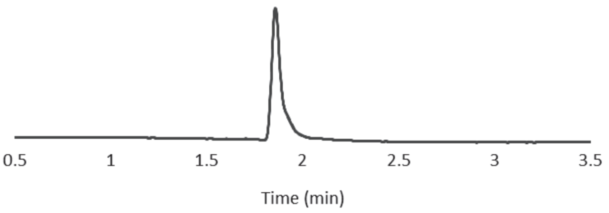


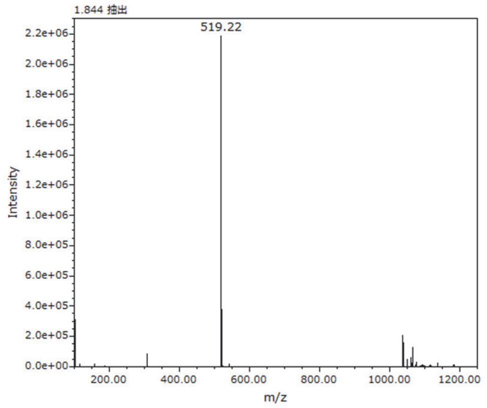


LC Chromatogram was monitored at 450 nm ($H_2O$-0.1% TFA/AcCN-0.1% TFA = 95/5 to 0/100, 3.5 min).
LRMS (ESI$^+$): $m/z$ = 519.2 $(M+H)^+$
